# Ligand stabilization enables physiological GITR signaling and antitumor immunity

**DOI:** 10.64898/2026.05.04.722444

**Authors:** Tesfahun Dessale Admasu, Keerthi Sadanala, Dnyaneshwar Kalyane, Hongqiang Wang, Seokyoung Yoon, Kim WooHyun, Madhusudhanarao Katiki, Jeremy Rudnick, Murali Ramachandran, John Yu

## Abstract

Agonistic antibodies targeting the costimulatory receptor GITR have failed in clinical trials despite a strong preclinical rationale, revealing fundamental limitations of receptor-targeted approaches to TNF superfamily agonism. Here, we demonstrate that pharmacological stabilization of the trimeric structure of human GITRL represents a mechanistically distinct strategy to overcome these limitations. Small-molecule stabilization of GITRL preserves its membrane residency. It enables bidirectional signaling between antigen-presenting cells and T cells, a physiological context that receptor-targeted antibodies bypass through non-physiological clustering of GITR. In humanized GITR/GITRL double-knock-in mice bearing syngeneic tumors, GITRL stabilization expands cytotoxic CD8⁺ T cells, selectively depletes intratumoral regulatory T cells, activates APCs, and restructures the immune spatial architecture. In patient-derived tumor explants, GITRL stabilization reactivates suppressed CD8⁺ T cells in an APC-dependent manner. Our findings establish ligand stabilization as a mechanistically distinct therapeutic strategy and provide a framework for engaging TNF receptor superfamily costimulatory pathways by modulating their native ligands.

**Teaser:** A small molecule stabilizes GITRL to activate tumor-fighting immune cells and reshape the tumor microenvironment.

## Introduction

Immune checkpoint inhibitors have transformed cancer treatment, yet durable responses remain limited in immunologically “cold” solid tumors such as glioblastoma, pancreatic adenocarcinoma, and treatment-refractory melanoma (*1, 2*). In these malignancies, regulatory T cells (Tregs) and tumor-associated macrophages (TAM) establish dominant suppressive networks that extinguish cytotoxic lymphocyte activity, prevent productive antigen presentation, and block formation of immune memory (*3, 4*). Effector CD8^+^ T cells become numerically depleted, metabolically constrained, and progressively exhausted, while myeloid cells fail to sustain the costimulatory signals required for T cell priming and effector differentiation (*5*). Therapeutic strategies capable of simultaneously reinforcing effector function and dismantling regulatory architecture are critically needed. The TNF receptor superfamily member GITR (glucocorticoid-induced TNFR-related protein; *TNFRSF18*) represents a compelling target because its engagement can enhance effector T cell proliferation, cytokine production, and resistance to Treg-mediated suppression (*6, 7*). Despite strong preclinical evidence, however, clinical trials of agonistic GITR antibodies have failed to demonstrate meaningful efficacy, even when combined with PD-1 or CTLA-4 blockade (*8–10*). This translational gap has revealed fundamental uncertainties regarding how TNF receptor superfamily costimulatory pathways should be therapeutically targeted. Structural and biochemical studies offer mechanistic insight into this failure. GITR signaling is initiated by its natural ligand, GITRL (*TNFSF18*), which is expressed as a membrane-bound homotrimer on antigen-presenting cells (APCs) (*11, 12*). The trimeric architecture of human GITRL, distinct from the dimeric structure of its murine orthologue, is essential for stable receptor clustering and sustained NF-κB activation (*11, 13*). Antibody agonists to GITR bypass the ligand, generating receptor oligomers that lack the geometric precision and cellular context of physiological GITR-GITRL engagement. Consequently, antibodies produce transient, incomplete signaling that fails to recapitulate the costimulatory synergy between GITR and canonical T cell activation pathways (*14*). Moreover, antibody-mediated receptor crosslinking does not engage APC-derived GITRL, thereby excluding bidirectional signaling interactions that may be required to coordinate T cell activation with APC maturation. Emerging evidence suggests that GITRL itself is not a passive ligand but can transduce reverse signals into APCs upon receptor binding (*15*). Reverse signaling through GITRL has been shown to activate NF-κB in macrophages and dendritic cells (DCs), upregulate costimulatory molecules, and promote inflammatory cytokine secretion (*16–18*). This raises the possibility that effective GITR agonism requires simultaneous forward signaling into T cells and reverse signaling into APCs, a dual-axis activation that receptor-targeted antibodies cannot deliver. Restoring this bidirectional signaling architecture may be essential for establishing durable antitumor immunity. We hypothesized that pharmacological stabilization of surface GITRL on APCs would recapitulate physiological GITR co-stimulation and enable therapeutic activity that is not attainable by receptor-directed approaches.

Here, we report stabilization of GITRL as a ligand-directed strategy for therapeutic GITR pathway agonism. We demonstrate that pharmacological preservation of GITRL surface expression and function maintains the cellular context of APC-T cell co-stimulation, enables bidirectional immune activation, and drives potent antitumor immunity in humanized mouse models. These findings establish ligand stabilization as a mechanistically distinct framework for targeting TNF receptor superfamily members and position GITRL as a tractable therapeutic node whose functional modulation offers advantages over conventional receptor-directed strategies.

## Results

### APC-restricted GITRL expression marks immune-competent tumor microenvironments

Bulk TCGA analyses showed elevated *TNFSF18* (GITRL) expression in tumors relative to matched normal tissues (Supplementary Fig. 1a). Further, analysis of public single-cell tumor atlases revealed that *TNFSF18* expression is highly enriched in myeloid APCs, with minimal expression detected in malignant cells or T cell populations (Supplementary Fig. 1b). Pan-cancer immune deconvolution demonstrated strong positive correlations between *TNFSF18* abundance and ImmuneScore, StromalScore, and IFN-γ-associated inflammatory signatures, indicating preferential enrichment in immune-infiltrated tumor microenvironments (TME) (Supplementary Fig. 1c). In lower-grade glioma, higher *TNFSF18* expression showed a trend toward improved survival in temozolomide-treated patients, whereas *TNFRSF18* (GITR) alone failed to stratify outcome (Supplementary Figs. 1d-e). In contrast, high co-expression of *TNFSF18* and *TNFRSF18* was associated with a survival benefit (Supplementary Fig. 1f), supporting the concept that ligand availability, rather than receptor expression alone, defines clinically relevant immune competence.

### GITRL stabilization enables bidirectional APC-T cell signaling

To interrogate the molecular requirements for GITRL signaling, we used KROS-101, a small molecule developed by Kairos Pharma, Ltd (Supplementary Fig. 2a). This small molecule selectively bound recombinant trimeric GITRL with nanomolar affinity (KD = 340 nM) (Supplementary Fig. 2b) with negligible affinity for the dimeric form. To provide structural insight into this selectivity, molecular docking analysis revealed that KROS-101 engages a central cavity formed exclusively (ΔG = −47 KCal/mol) at the convergence of the three GITRL protomers (Supplementary Fig. 2c-d), a binding site that is geometrically absent in the dimeric configuration. This structural interpretation is consistent with the binding data and supports the notion that KROS-101 acts as a trimer-selective stabilizer rather than a non-specific ligand. In primary APC monocultures derived from healthy donor PBMCs, KROS-101 stabilized surface GITRL and prolonged membrane residence on both DCs and M1 macrophages (Fig. 1a, b) without affecting MHC-II or CD86 (Supplementary Fig. 2e-f). Surface GITRL rose transiently and then declined progressively in vehicle-treated cultures but remained at or near peak levels with KROS-101, consistent with prolonged stabilization of the surface GITRL pool. Together, these findings support a model in which sustained membrane presentation of trimeric GITRL promotes productive GITRL-GITR engagement by maintaining a geometrically constrained ligand configuration that favors multivalent GITR clustering and signal amplification. To confirm that stabilization preserves physiological signaling, we performed NF-κB luciferase reporter assays in Jurkat cells. Trimeric rhGITRL induced robust NF-κB activation, whereas monomeric rhGITRL failed to recapitulate this response (Fig. 1c). KROS-101 alone showed no significant activity, whereas preincubation with trimeric rhGITRL fully preserved NF-κB activation (Fig. 1d). Notably, KROS-101 did not further increase NF-κB activation at saturating trimeric rhGITRL but significantly enhanced signaling at a sub-saturating concentration to levels comparable to saturating ligand (Fig. 1d), indicating enhanced GITRL signaling under ligand-limiting conditions. To assess the functional consequences of membrane ligand stabilization, we established defined tri-culture systems comprised of purified human CD8⁺ T cells, autologous Tregs, and APCs. In APC-T cell co-cultures, KROS-101 treatment induced robust bidirectional activation. APCs upregulated MHC-II and CD86 on both macrophages and DCs (Fig. 1e, f) while CD8⁺ T cells showed increased CD25 expression (Fig. 1g), and enhanced intracellular IFN-γ and TNF-α accumulation (Fig. 1h, i). Concurrently, Treg cells exhibited reduced FoxP3 expression and diminished IL-10 and TGF-β production (Fig. 1j-l), consistent with destabilization of suppressive function. In contrast, in the absence of APCs, Treg cells induced a dysfunctional phenotype in CD8⁺ T cells characterized by increased PD-1 and LAG-3 expression, whereas KROS-101 had no measurable effect (Fig. 1m). Thus, the suppressive-Treg reversal observed in Fig. 1j-l requires APC-presented GITRL and is not expected in CD8:Treg cultures lacking the ligand-bearing APC compartment.

**Figure 1:**
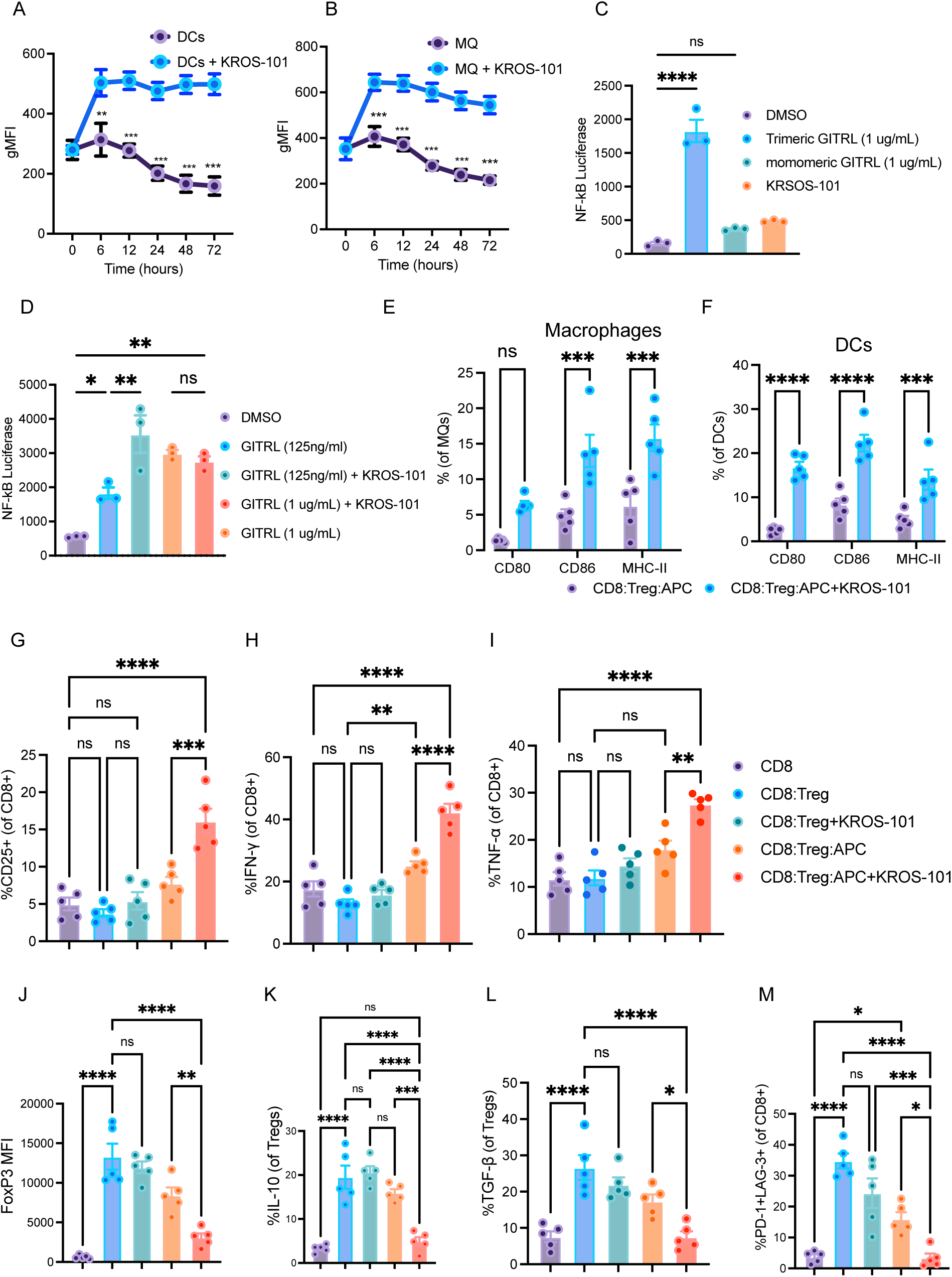
KROS-101 stabilizes GITRL and enhances APC-T cell signaling. **a,b**, Surface GITRL expression on DCs (**a**) and MQs (**b**) over time following treatment with KROS-101, showing prolonged ligand surface stability. **c**, NF-κB luciferase reporter activity in Jurkat cells stimulated with trimeric GITRL, monomeric GITRL, or KROS-101 alone. Trimeric GITRL induced robust NF-κB activation, whereas monomeric GITRL and KROS-101 alone showed minimal activity. **d**, NF-κB reporter activity following stimulation with trimeric GITRL, demonstrating that KROS-101 did not further increase NF-κB activation at a saturating GITRL concentration but significantly enhanced signaling at a sub-saturating concentration. **e,f**, Expression of antigen-presentation and co-stimulatory molecules (CD80, CD86, MHCII) on MQs (**e**) and DCs (**f**) in CD8:Treg:APC co-cultures with or without KROS-101. **g-i**, Activation of CD8⁺ T cells in co-culture systems containing CD8 T cells alone or combined with Tregs and APCs. KROS-101 increased CD25 expression (**g**) and enhanced cytokine production, including IFNγ (**h**) and TNFα (**i**), in the presence of APCs. **j-l**, Effects of KROS-101 on Treg phenotype and suppressive cytokines. KROS-101 reduced FoxP3 expression (**j**) and decreased IL-10 (**k**) and TGF-β (**l**) production by Tregs. **m**, Frequency of exhausted PD-1⁺LAG-3⁺ CD8 T cells across co-culture conditions, showing reduced exhaustion following KROS-101 treatment in APC-containing systems. These effects were observed in APC-containing tri-cultures (CD8:Treg:APC) but not in the absence of APCs (CD8:Treg cultures lacking GITRL-bearing APCs). Data are mean ± s.e.m. Each dot represents an independent donor. Statistical comparisons were performed using one-way ANOVA with Tukey’s test for multiple comparisons. ns, not significant; ** p < 0.01; *** p < 0.001; **** p < 0.0001.

### Ligand availability governs responsiveness to GITRL stabilization in human TILs

To determine whether endogenous GITRL availability constrains the activity of KROS-101 in human tumors, we first profiled GITRL expression across immune subsets within freshly isolated TILs from glioma, glioblastoma (GBM), triple-negative breast cancer, ependymoma, and uterine carcinoma specimens (Fig. 2a, b). GITRL expression varied markedly across donors and tumor types but was consistently enriched on intratumoral DCs and macrophages, establishing substantial inter-patient heterogeneity in ligand availability (Fig. 2b, c). These data indicate that APC GITRL abundance is a candidate response biomarker rather than a uniform property of all tumors. We next asked whether this heterogeneity was translated into differential functional responsiveness. In TIL cultures derived from high GITRL-expressing tumors, GITRL stabilization enhanced autologous tumor cell killing (Fig. 2e, f) and increased production of IFN-γ, TNF-α, and IL-2 (Fig. 2d, g). In contrast, TILs from low GITRL-expressing tumors showed minimal cytotoxic response (Fig. 2h, i). Across donors and cancer types, baseline GITRL abundance on macrophages and DCs correlated with the magnitude of stabilization-induced cytotoxicity (Fig. 2k-m). To confirm that APC-expressed GITRL is required for this effect, we disrupted ligand engagement using a GITRL neutralizing antibody during autologous killing assays. GITRL blockade abolished the enhancement of cytotoxicity conferred by GITRL stabilization (Fig. 2j). Together, these data identify baseline GITRL abundance on tumor-resident APCs as a quantitative determinant of responsiveness to GITRL stabilization and nominate it as a candidate predictive biomarker to guide patient selection, with tumors exhibiting low APC GITRL expression predicted to derive limited benefit from monotherapy.

**Figure 2:**
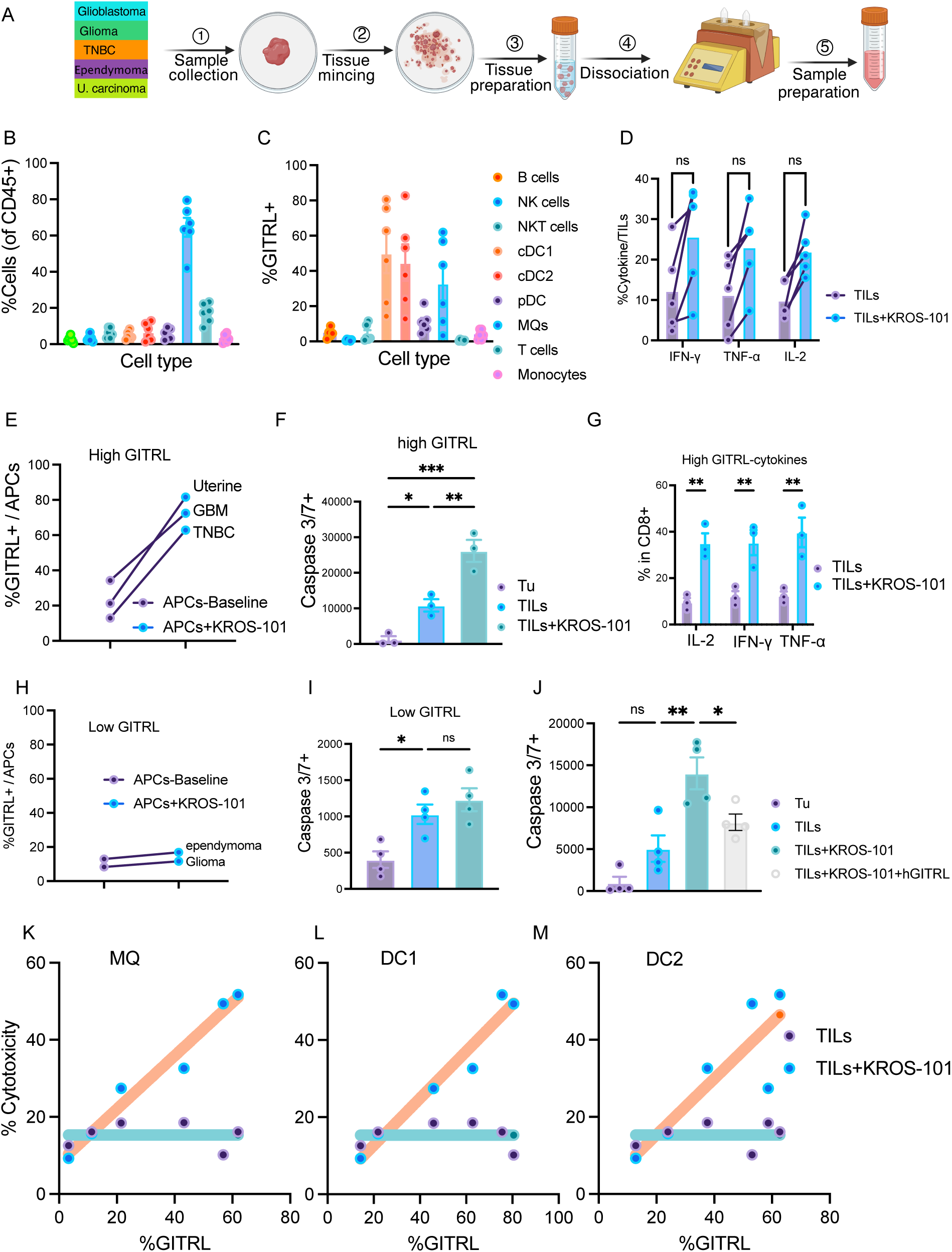
KROS-101 enhances TIL-mediated tumor cell killing in human tumors with high APC GITRL expression. **a,** Workflow for isolation and functional analysis of tumor-infiltrating lymphocytes (TILs) from freshly resected human tumors, including glioblastoma (GBM), glioma, ependymoma, triple-negative breast cancer (TNBC) brain metastasis, and uterine carcinoma brain metastasis. Tissue was minced, dissociated, and processed to obtain single-cell suspensions for immune profiling and functional assays. **b**, Composition of tumor-infiltrating immune cells among CD45⁺ populations across patient samples. **c**, Frequency of GITRL⁺ cells among tumor-infiltrating immune populations, showing enrichment in APCs, particularly DCs and MQs. **d**, Cytokine production by TILs following short-term stimulation in the presence or absence of KROS-101. **e**, Frequency of GITRL⁺ APCs in tumors with high baseline GITRL expression (GBM, TNBC, uterine carcinoma) following *ex vivo* treatment with KROS-101. **f**, Tumor cell apoptosis measured by caspase-3/7 activation following co-culture with TILs in high-GITRL tumors, demonstrating enhanced tumor cell killing in the presence of KROS-101. **g**, Cytokine production (IL-2, IFNγ, TNFα) by CD8⁺ TILs from high-GITRL tumors following KROS-101 treatment. **h**, Frequency of GITRL⁺ APCs in tumors with low baseline GITRL expression (glioma and ependymoma) following KROS-101 exposure. **i**, Tumor cell apoptosis following TIL co-culture in low-GITRL tumors. KROS-101 produced minimal enhancement of cytotoxicity in this context. **j**, Tumor cell apoptosis measured by caspase-3/7 activation following co-culture with TILs in high-GITRL tumors in the presence of GITRL neutralizing antibody, demonstrating GITRL blockade abolished the enhancement of cytotoxicity conferred by GITRL stabilization. **k-m**, Correlation between APC GITRL expression and TIL-mediated tumor cytotoxicity for MQs (**k**), cDC1 (**l**), and cDC2 (**m**), demonstrating that higher APC GITRL levels predict stronger KROS-101-dependent tumor cell killing. Statistical comparisons were performed using a t-test or one-way ANOVA. ns, not significant; **p < 0.01; ***p < 0.001. Each dot represents an individual patient sample. The experimental design in panel A was created in BioRender. Admasu, T. (2026) https://BioRender.com/n0ofqlw.

### GITRL stabilization drives antitumor immunity in hGITR/hGITRL humanized mice

Because murine GITRL forms a dimer that is not recognized by KROS-101, conventional syngeneic models were not suitable. We therefore performed all *in vivo* studies in hGITR/hGITRL double-humanized knock-in mice that recapitulate human GITR-GITRL interactions (Fig. 3a). In this mouse model, the extracellular domains of endogenous GITR and GITRL are replaced with their human counterparts while preserving intact murine intracellular signaling and immune architecture. To compare ligand stabilization directly with antibody-mediated GITR agonism, we evaluated KROS-101 and clinical-stage GITR agonist antibody TRX518 head-to-head in checkpoint-resistant B16-F10-Luc2 and checkpoint-responsive MC38 tumors. In B16-F10-Luc2, KROS-101 significantly reduced tumor growth relative to vehicle, whereas TRX518 did not; neither agent significantly prolonged survival (Fig. 3b, c). Notably, a substantial proportion of KROS-101-treated mice reached humane endpoints because of dry tumor ulceration rather than progressive tumor growth. In MC38, both KROS-101 and TRX518 significantly reduced tumor growth and prolonged survival relative to vehicle (Fig. 3d, e). KROS-101 also reduced tumor progression and improved survival in YUMM1.7 melanoma (Fig. 3f, g) and significantly prolonged survival in orthotopic GL261 glioma (Fig. 3h).

**Figure 3:**
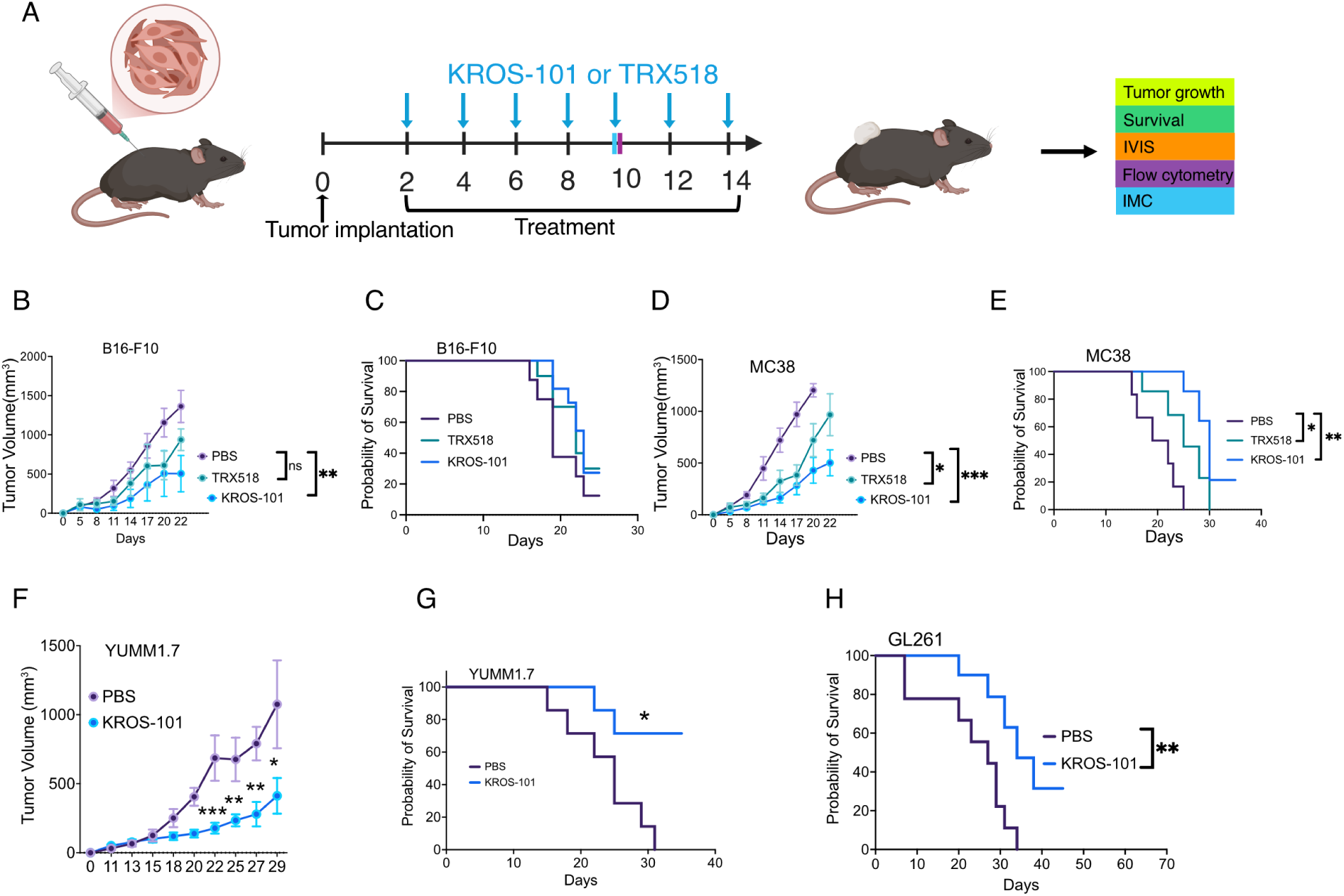
KROS-101 suppresses tumor growth and improves survival in hGITR/hGITRL double-knock-in syngeneic mouse tumor models. **a**, Experimental design for *in vivo* efficacy studies. Mice bearing established tumors were treated with KROS-101 or TRX518 beginning on day 2 after tumor implantation. Tumor growth and survival (in B16-F10-Luc2, MC38, and GL261) and immune profiling by flow cytometry (in B16-F10 and MC38) and imaging mass cytometry (IMC, in B16-F10) were performed at the indicated time points. **b,c**, Tumor growth kinetics (**b**) and Kaplan-Meier survival curves (**c**) in the B16-F10 melanoma model following treatment with TRX518 or KROS-101. KROS-101 significantly reduced tumor growth, n = 8 mice per group. **d,e**, Tumor volume (**d**) and survival (**e**) of mice bearing MC38 colon carcinoma tumors treated with TRX518 or KROS-101, n = 7 mice per group. **f**,**g**, Tumor volume (**f**) and survival (**g**) of mice bearing YUMM1.7 melanoma tumors treated with PBS or KROS-101, n = 8 mice per group. **h**, Kaplan-Meier survival curves in GL261 glioblastoma-bearing mice treated with PBS or KROS-101, n = 8 mice per group. Survival curves were compared using the log-rank (Mantel-Cox) test. Tumor volumes are shown as mean ± s.e.m. * p < 0.05; ** p < 0.01. The experimental design in panel A was created in BioRender. Admasu, T. (2026) https://BioRender.com/9q24y17)

### GITRL stabilization enhances effector T cell responses while depleting Tregs

To define the immunologic impact of KROS-101 and directly benchmark it against the clinical-stage GITR antibody agonist TRX518, we performed high-dimensional flow cytometric immunoprofiling of TILs in MC38 and B16-F10 tumor-bearing mice at day 7 post-treatment initiation (Supplementary Fig. 3a, b). In MC38 tumors, KROS-101 increased total CD45+ immune infiltration and expanded intratumoral CD8+ T cells while reducing Tregs, resulting in an increased CD8/Treg ratio; importantly, KROS-101 produced significantly greater increases in TILs, CD8+ T cells, and the CD8/Treg ratio than TRX518 (Fig. 4a-f). This expansion was accompanied by enrichment of CD44+CD62L− effector CD8+ T cells which was significantly greater with KROS-101 than with TRX518 (Fig. 4b), and increased GITR, TNF-α, IFN-γ, and granzyme B expression, with significantly greater GITR and granzyme B responses than TRX518 (Fig. 4c). These cells also showed a significant increase in GITR expression in MC38 tumors (Fig. 4c, g). Examination of exhaustion markers revealed that KROS-101 reduced PD-1+TIM-3+ CD8+ cells, with a significant difference compared with TRX518 (Fig. 4d). Further, KROS-101 selectively depleted intratumoral Treg cells (CD4+Foxp3+) (Fig. 4a, f). Residual Tregs within KROS-101-treated tumors displayed reduced expression of the suppressive markers LAP and IL-10, both significantly lower than with TRX518 (Fig. 4h). Given that KROS-101 targets GITRL expression by APCs, we next assessed tumor-associated myeloid populations. KROS-101 increased the M1-like (CD86⁺MHCII^hi^) macrophage compartment without significantly altering cDC1, cDC2, or total macrophage frequencies in MC38 tumors (Fig. 4i). KROS-101 also increased GITRL expression on cDC1s (Fig. 4j) and cDC1-associated PD-L1, CD86, and MHC-II expression (Fig. 4k), consistent with enhanced APC activation.

**Figure 4:**
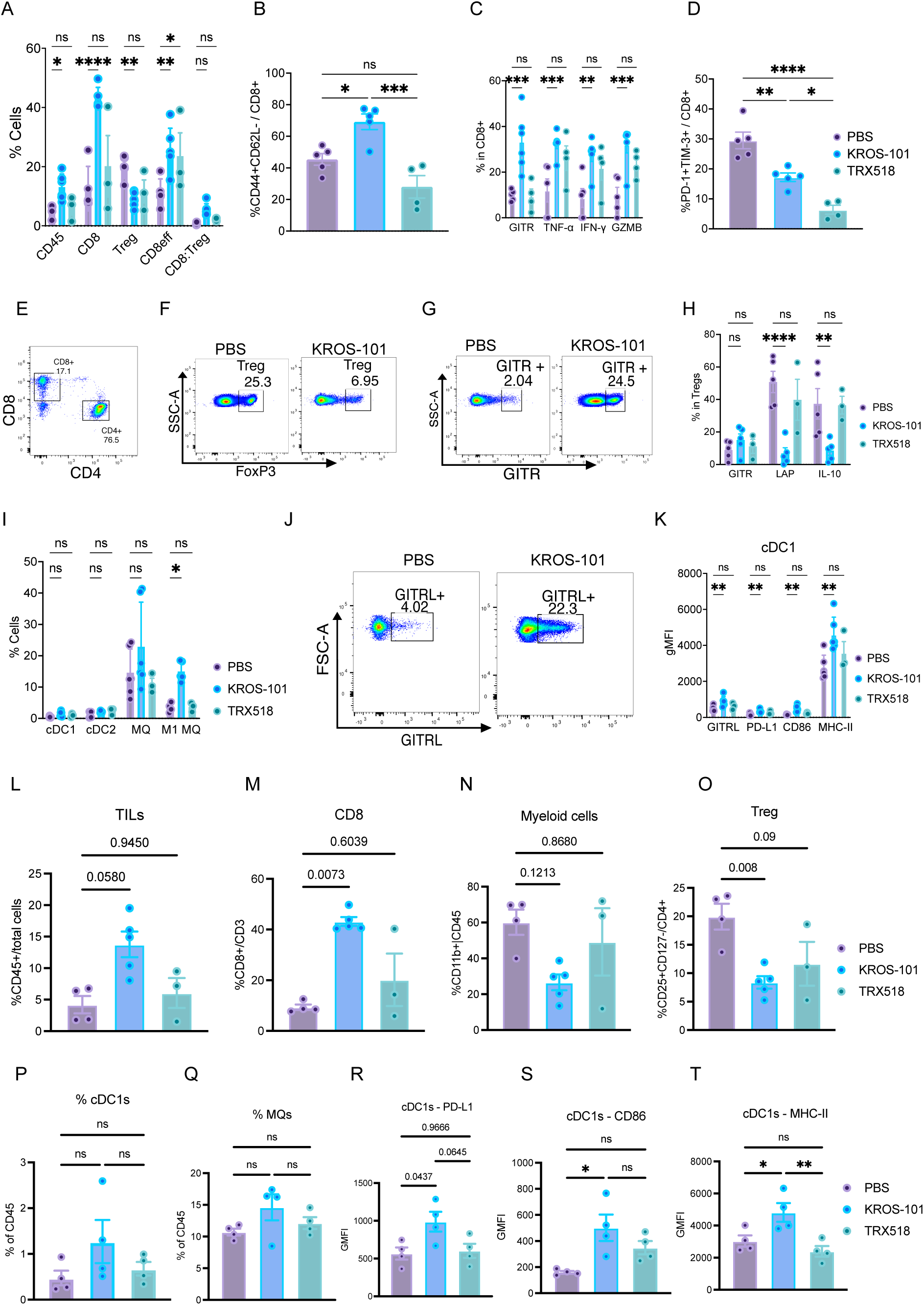
KROS-101 remodels the tumor immune microenvironment by enhancing effector CD8⁺ T cell responses and reducing Treg-mediated suppression. **a-k**, Immune profiling of MC38 tumors following treatment with PBS, KROS-101, or TRX518. (**a**) Frequencies of CD45+ TILs, CD8+ T cells, Tregs, CD8+ effector T cells, and CD8:Treg ratio. (**b**) Frequency of CD44+CD62L− effector CD8+ T cells. (**c**) GITR, TNF-α, IFN-γ, and granzyme B expression in CD8+ T cells. (**d**) Frequency of PD-1+TIM-3+ CD8+ T cells. (**e-g**) Representative flow cytometry plots showing CD4+ and CD8+ T cells (**e**) Tregs (**f**) and GITR+ CD8+ T cells (**g**). (**h**) GITR, LAP, and IL-10 expression in Tregs. (**i**) Frequencies of cDC1, cDC2, macrophages (MQ), and M1-like macrophages. (**j**) Representative flow cytometry plots showing GITRL expression on cDC1s. (**k**) GITRL, PD-L1, CD86, and MHC-II expression on cDC1s. **l-t**, Immune profiling of B16-F10 tumors following treatment with PBS, KROS-101, or TRX518. (**l**) Frequency of CD45+ TILs. (**m**) Frequency of CD8+ T cells within the CD3+ compartment. (**n**) Frequency of CD11b+ myeloid cells within CD45+ cells. (**o**) Frequency of Tregs (CD25+CD127−/CD4+). (**p**) Frequency of cDC1s within CD45+ cells. (**q**) Frequency of macrophages within CD45+ cells. **r-t**, Expression of PD-L1 (**r**) CD86 (**s**) and MHC-II (**t**) on cDC1s. Data are presented as mean ± s.e.m. Statistical comparisons were performed using one-way ANOVA with Tukey’s test. *P < 0.05; **P < 0.01; ***P < 0.001; ****P < 0.0001; ns, not significant. Each dot represents one biological replicate (individual mouse).

We next assessed whether this immunologic remodeling extended to the poorly immunogenic B16-F10 model. KROS-101 increased total CD45+ immune infiltration and CD8+ T cells while reducing intratumoral Tregs, with significantly greater TIL infiltration and Treg depletion than TRX518 (Fig. 4l-o). Total myeloid populations, neutrophils, MDSC, cDC1, or macrophage frequencies were not significantly altered (Fig. 4p, q and Supplementary Fig. 4e), whereas KROS-101 increased cDC1-associated PD-L1, CD86, and MHC-II expression (Fig. 4r-t). Additional profiling of B16-F10 tumors showed increased IFN-γ and granzyme B production and reduced TIGIT and TIM-3 expression in CD8+ T cells (Supplementary Fig. 4c), together with Treg depletion and reduced LAP expression (Supplementary Fig. 4d). These effects were tumor-restricted, as blood and splenic Treg and effector T cell frequencies remained unchanged (Supplementary Fig. 4a, b). Collectively, these findings demonstrate that KROS-101 produces consistent remodeling of the tumor immune microenvironment across MC38 and B16-F10 tumors by enhancing effector T cell responses, depleting Tregs, and activating APCs, with multiple effects significantly greater than those achieved by antibody-mediated GITR agonism.

### GITRL stabilization promotes APC-CD8 T cell interactions within tumor parenchyma

To interrogate the spatial organization underlying KROS-101’s therapeutic activity, we performed H&E histology, CD3 immunohistochemistry (IHC), and imaging mass cytometry (IMC) on B16-F10 tumor sections. H&E staining revealed markedly increased lymphocytic infiltration in KROS-101-treated tumors relative to vehicle controls (Supplementary Fig. 5a, b). CD3 IHC confirmed a substantial increase in T cell infiltration, with KROS-101-treated tumors showing significantly higher CD3⁺ cell counts across all three quantification metrics, total CD3⁺ cells, CD3⁺ cells per gram of tumor, and CD3⁺ cells per mm², compared to PBS-treated controls (Supplementary Fig. 5c-g). IMC further resolved the spatial architecture of this response at single-cell resolution. Vehicle-treated tumors exhibited predominant peripheral localization of CD8⁺ T cells with limited intratumoral penetration (Fig. 5a), whereas KROS-101 treatment substantially increased CD8⁺ T cell density within tumor parenchyma (Fig. 5b, c) and their representation among CD45⁺ infiltrating leukocytes (Fig. 5d). Quantification of CD4⁺ T cells in the same IMC dataset (Supplementary Fig. 5h and i) showed only modest changes with KROS-101, in contrast to the marked increase in CD8⁺ T cells, indicating that GITRL stabilization primarily expands the intratumoral cytotoxic CD8⁺ compartment. Spatial neighborhood analysis revealed that CD8⁺ T cells were significantly enriched within a 30 μm radius of APCs in KROS-101-treated tumors, and the majority of these APC-proximal CD8⁺ T cells co-expressed granzyme B, indicating active cytotoxic programming at sites of antigen encounter (Fig. 5e, f). Myeloid compartment analysis revealed spatial reprogramming without changes in overall abundance. Although total F4/80⁺ TAM density was unchanged (Supplementary Fig. 6a), KROS-101 significantly increased the frequency of MHC-II⁺CD80⁺ inflammatory TAMs within tumor nests (Supplementary Fig. 6b), and these activated macrophages were observed in close spatial proximity to CD8⁺ T cell aggregates (Fig. 5a, b). In parallel, total CD11c⁺ DC numbers were significantly elevated in KROS-101-treated tumors (Supplementary Fig. 6c), and CD11c⁺MHC-II⁺ DCs localized to inflamed tumor regions (Fig. 5a, b, g). F4/80⁺CD11c⁺ cells were more abundant and spatially enriched near CD8⁺ T cell aggregates in KROS-101-treated tissue relative to vehicle controls (Fig. 5h), consistent with a role for antigen-presenting myeloid cells in sustaining local cytotoxic effector activity within these immune niches.

**Figure 5:**
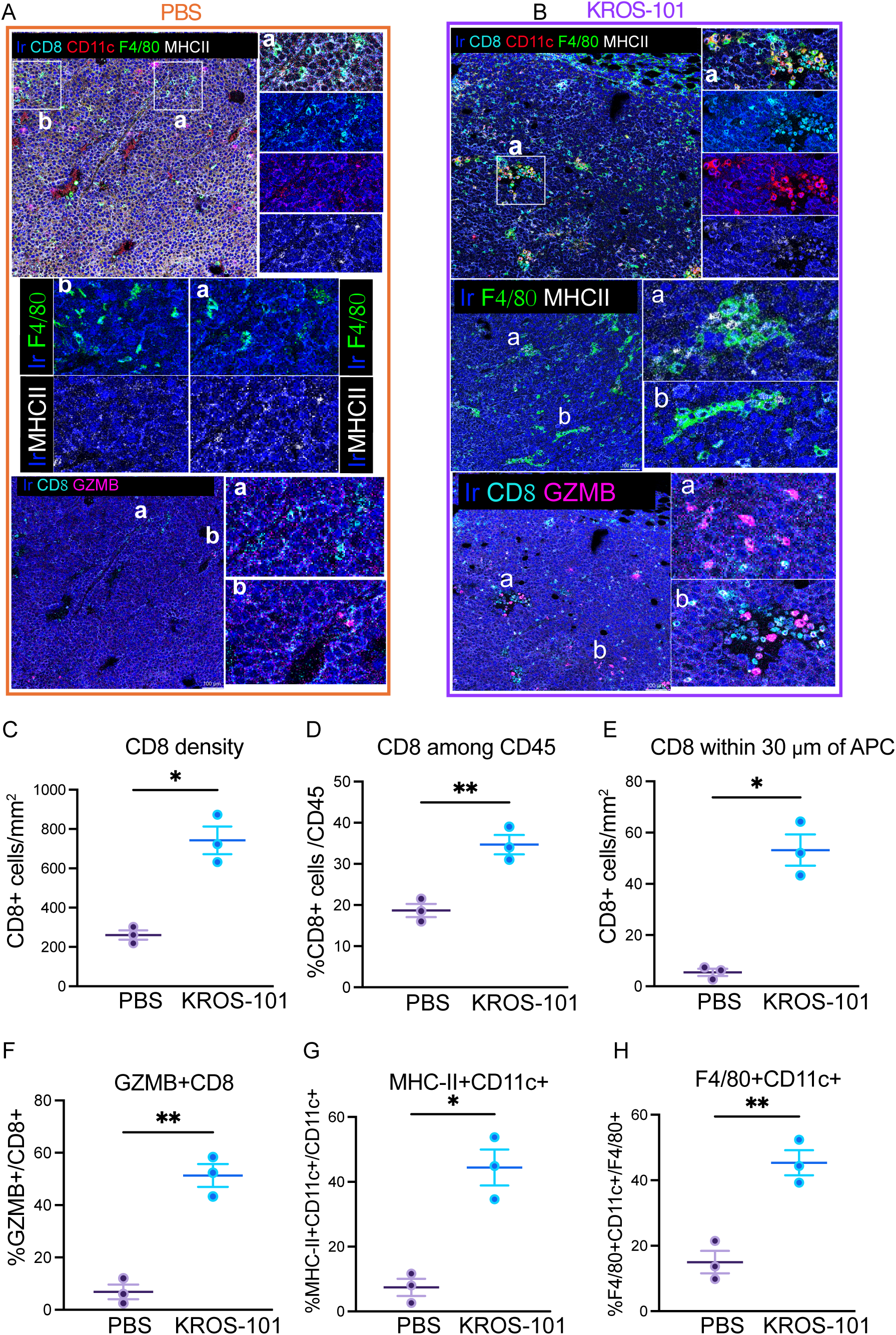
KROS-101 promotes spatial organization of APC-CD8 interactions and enhances intratumoral cytotoxic T cell activity. **a,b**, Representative imaging mass cytometry (IMC) of tumor sections from PBS-treated (**a**) and KROS-101 treated (**b**) mice stained for iridium (Ir, nuclear), CD8 (cyan), CD11c (red), F4/80 (green), and MHCII (white). Insets show regions enriched for APCs and CD8⁺ T cells. Additional overlays highlight F4/80⁺ macrophages and MHCII⁺ APCs, as well as granzyme B (GZMB) expression in CD8⁺ T cells. Scale bars, 100 µm. **c**, Quantification of intratumoral CD8⁺ T cell density (cells/mm^2^) showing increased CD8 infiltration following KROS-101 treatment. **d**, Frequency of CD8⁺ T cells among CD45⁺ leukocytes in tumors. **e**, Spatial proximity analysis showing the proportion of CD8⁺ T cells located within 30 μm of APCs, indicating enhanced APC-T cell interactions in KROS-101 treated tumors. **f**, Frequency of cytotoxic GZMB⁺ CD8⁺ T cells in tumors following KROS-101 treatment. **g**, Density of activated antigen-presenting cells (MHCII⁺CD11c⁺) within tumors. **h**, Frequency of F4/80⁺CD11c⁺ macrophage-like APCs among tumor-associated macrophages. Statistical comparisons were performed using an unpaired t-test. * p < 0.05; ** p < 0.01. Data shown as mean ± s.e.m.

### GITRL stabilization drives MHC-restricted, antigen-specific cytotoxic memory

To determine whether the immune remodeling induced by KROS-101 *in vivo* translated into functional cytotoxic competence, we evaluated the *ex vivo* killing capacity of tumor-infiltrating CD8⁺ T cells. CD8⁺ TILs from KROS-101-treated tumors exhibited markedly enhanced tumor cell killing *ex vivo* compared with CD8⁺ TILs from vehicle-treated controls in both B16-F10-Luc2 and MC38 models (Fig. 6a, b). This increased cytotoxic activity was accompanied by elevated IFN-γ and increased GZMB accumulation (Fig. 6c). To define the effector mechanisms underlying this enhanced cytotoxicity, we interrogated the contributions of cytokine signaling, antigen presentation, and cytotoxic execution pathways. First, we showed that neutralization of IFN-γ partially reduced KROS-101-induced tumor cell killing (Fig. 6d). Inhibition of perforin-dependent granule acidification with concanamycin A also suppressed tumor cell death (Fig. 6e). Finally, MHC class I/II-blocking antibodies nearly abolished tumor cell killing by CD8⁺ TILs, demonstrating that cytotoxicity was dependent on MHC-restricted antigen presentation (Fig. 6f). Lastly, to evaluate tumor antigen-specific recall responses, splenocytes were isolated from vehicle-and KROS-101-treated mice. Upon *ex vivo* restimulation with MC38 lysate-pulsed antigen-presenting cells, splenocytes from KROS-101-treated mice exhibited increased IFN-γ accumulation compared with vehicle controls (Supplementary Fig. 6d). This recall response was abrogated by MHC class I blockade during restimulation (Supplementary Fig. 6e), confirming that KROS-101 promotes antigen-experienced CD8⁺ T cells that mount MHC-I restricted effector recall responses. However, because these recall assays measured CD8⁺ T cells and MHC-I blockade, they do not exclude additional MHC-II-restricted effects on antigen-experienced CD4⁺ T cells.

**Figure 6:**
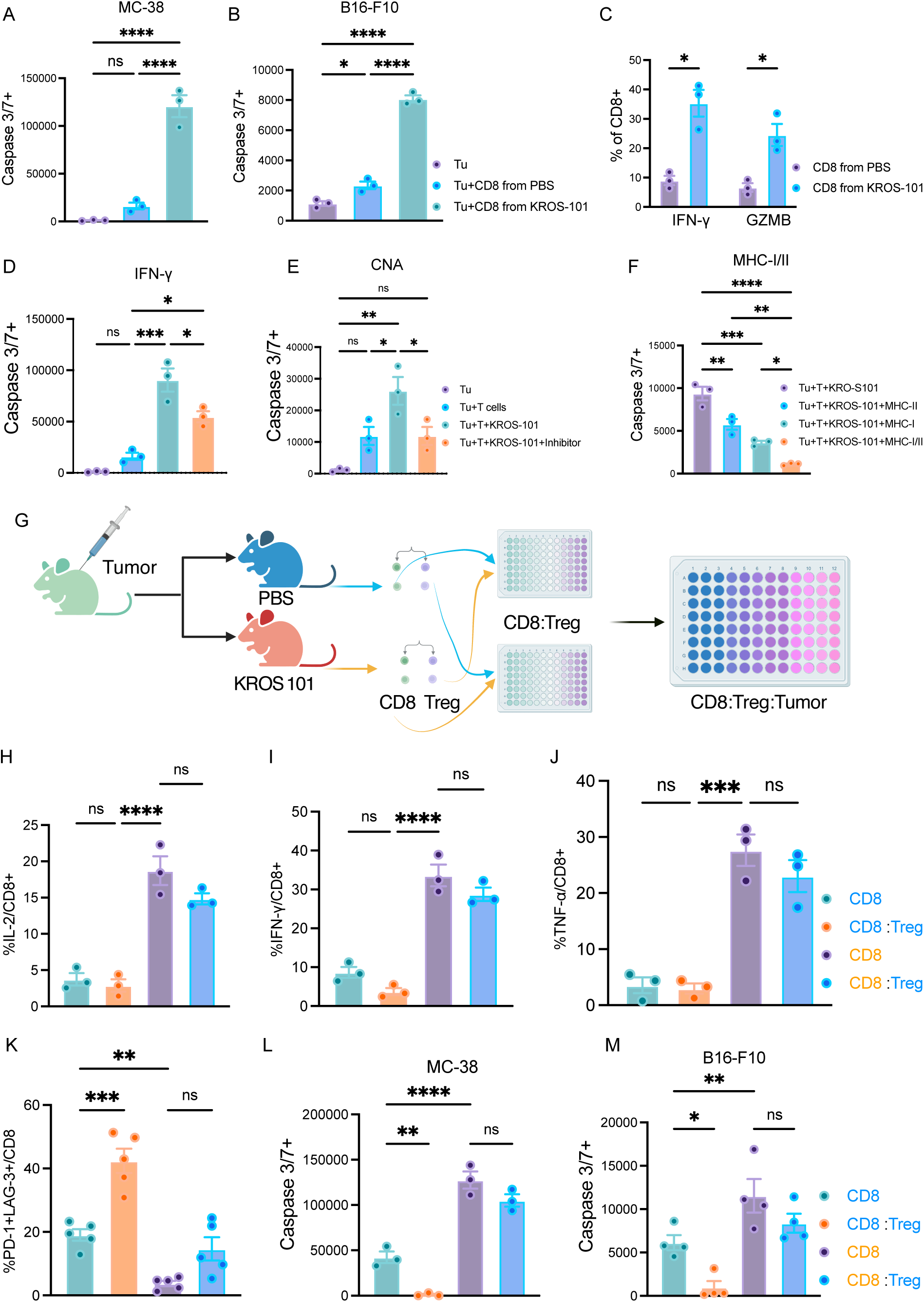
KROS-101 promotes MHC-restricted cytotoxicity and overcomes Treg-mediated suppression. **a,b**, Tumor cell killing measured by caspase-3/7 activation in MC38 (**a**) and B16-F10 (**b**) tumor cells co-cultured with CD8⁺ T cells isolated from PBS-or KROS-101-treated mice. CD8⁺ T cells from KROS-101 treated mice exhibited enhanced cytotoxic activity. **c**, Effector function of CD8⁺ T cells isolated from treated mice showing increased IFN-γ and GZMB production following KROS-101 treatment. **d**, Tumor cell killing in co-cultures treated with IFN-γ neutralizing antibody, demonstrating partial dependence of KROS-101-mediated cytotoxicity on IFN-γ signaling. **e**, Effects of perforin-mediated cytotoxicity inhibition using concanamycin A (CNA). Inhibition reduced tumor cell killing, indicating involvement of granule-mediated cytotoxic mechanisms. **f**, Blocking of MHC class I and/or class II during tumor cell killing assays shows that KROS-101 enhanced cytotoxicity depends on antigen presentation through MHC pathways. **g**, Experimental schematic of the *ex vivo* co-culture system in which CD8⁺ T cells and Tregs isolated from PBS-or KROS-101-treated mice are combined before co-culture with tumor cells. **h-j**, Cytokine production by CD8⁺ T cells in CD8-only or CD8:Treg co-cultures, showing increased IL-2 (**h**), IFN-γ (**i**), and TNF-α (**j**) by CD8⁺ T cells isolated from KROS-101-treated mice. **k**, Frequency of exhausted PD-1⁺LAG-3⁺ CD8⁺ T cells in co-culture conditions, demonstrating reduced exhaustion following KROS-101 treatment. **l,m**, Tumor cell killing in MC38 (**l**) and B16-F10 (**m**) tumor models in CD8-only or CD8:Treg co-cultures. KROS-101 enhanced CD8⁺ T cell-mediated cytotoxicity and overcame Treg-mediated suppression. Statistical comparisons by one-way ANOVA with Tukey’s test. ns, not significant; * p < 0.05; ** p < 0.01; *** p < 0.001; **** p < 0.0001. Data shown as mean ± s.e.m. Each dot represents an independent mouse. The experimental design in panel G was created in BioRender. Admasu, T. (2026) https://BioRender.com/vr61lva

### GITRL stabilization durably reprograms effector and regulatory T cells

To assess whether KROS-101-induced immune changes persist beyond acute treatment, we performed ex vivo co-culture assays using CD8⁺ T cells and Tregs isolated independently from PBS-or KROS-101-treated tumors (Fig. 6g). CD8⁺ T cells from KROS-101-treated tumors produced significantly higher levels of IL-2, TNF, and IFN-γ (Fig. 6h-j) and exhibited greater cytotoxic activity against both MC38 (Fig. 6l) and B16-F10 (Fig. 6m) target cells compared to CD8⁺ T cells from PBS-treated tumors. Most importantly, this functional superiority was maintained even when KROS-101-derived CD8⁺ T cells were co-cultured with Tregs from PBS-treated tumors, demonstrating resistance to extrinsic suppression. By contrast, PBS-derived CD8⁺ T cells showed impaired cytokine secretion (Fig. 6h-j) and cytotoxicity (Fig. 6l, m) under basal conditions, which was further reduced and associated with significantly elevated exhaustion (PD-1^+^LAG-3^+^) upon co-culture with PBS-derived Tregs (Fig. 6k). PBS-derived Tregs had no significant effect on the cytokine output (Fig. 6h-j), cytotoxicity (Fig. 6l, m), or exhaustion (Fig. 6k) state of KROS-101-derived CD8⁺ T cells. Together, these data demonstrate that KROS-101 induces durable functional reprogramming of CD8⁺ T cells that is intrinsically maintained and refractory to Treg-mediated suppression.

### Checkpoint blockade enhances the antitumor activity of KROS-101

To test whether checkpoint blockade complements GITRL stabilization, we evaluated KROS-101 in combination with anti-PD-1 or anti-CTLA-4 across MC38 and B16-F10 tumors (Fig. 7). In MC38, KROS-101 plus anti-PD-1 improved tumor growth control, although the difference from KROS-101 monotherapy did not reach statistical significance (Fig. 7a). The combination did not further improve tumor growth control over KROS-101 monotherapy, although it significantly prolonged survival relative to vehicle (Fig. 7b). In contrast, KROS-101 plus anti-CTLA-4 significantly improved tumor growth control relative to the single-agent treatments and produced greater survival benefit, including a significant improvement over KROS-101 and CTLA-4 monotherapy (Fig. 7c, d). In the poorly immunogenic B16-F10 model, KROS-101 plus anti-PD-1 significantly improved tumor growth control compared with single-agent treatments (Fig. 7e). This enhanced tumor control was accompanied by a survival benefit: neither KROS-101 nor anti-PD-1 monotherapy significantly prolonged survival relative to vehicle, whereas the combination did (Fig. 7f). Together, these findings demonstrate that checkpoint blockade enhances the antitumor activity of KROS-101, with the magnitude of benefit varying by tumor model and checkpoint target.

**Figure 7.**
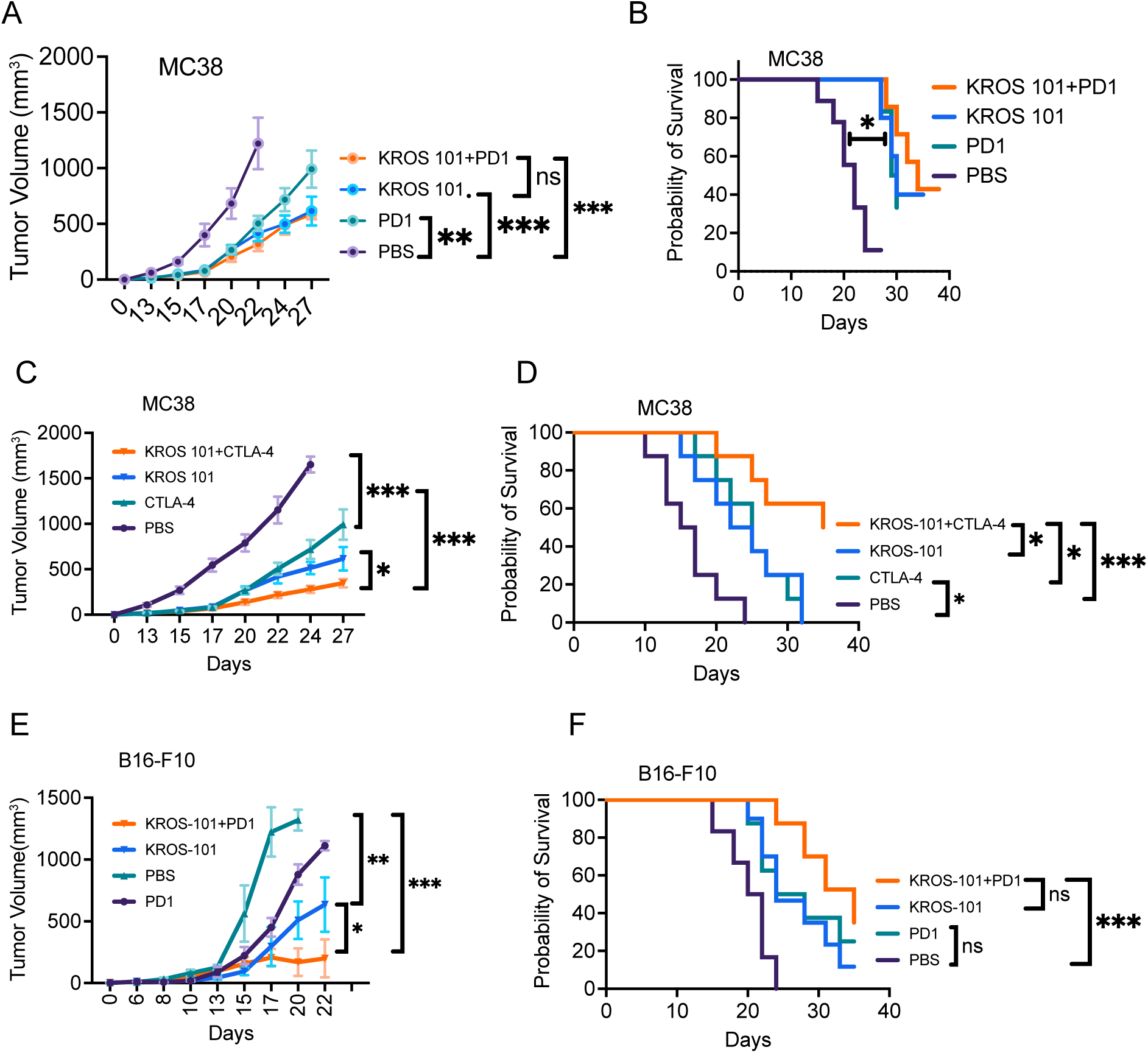
Checkpoint blockade enhances the antitumor activity of KROS-101. **a, b**, Antitumor activity of KROS-101 in combination with anti-PD-1 in MC38 tumor-bearing mice. Mice were treated with PBS, KROS-101, anti-PD-1 (clone RMP1-14, Bio X Cell; 200 µg/mouse, i.p.) or KROS-101 plus anti-PD-1. Tumor growth (**a**) and Kaplan-Meier survival (**b**) are shown (n = 8 mice per group). **c, d**, Antitumor activity of KROS-101 in combination with anti-CTLA-4 in MC38 tumor-bearing mice. Mice were treated with PBS, KROS-101, anti-CTLA-4 (clone 9H10, Bio X Cell; 100 µg/mouse, i.p.), or KROS-101 plus anti-CTLA-4. Tumor growth (**c**) and Kaplan-Meier survival (**d**) are shown (n = 8 mice per group). **e, f**, Antitumor activity of KROS-101 in combination with anti-PD-1 in B16-F10 tumor-bearing mice. Mice were treated with PBS, KROS-101, anti-PD-1 (clone RMP1-14, Bio X Cell; 200 µg/mouse, i.p.), or KROS-101 plus anti-PD-1. Tumor growth (**e,** n = 4 mice per group) and Kaplan-Meier survival (**f,** n = 7 mice per group) are shown. The combination significantly improved tumor growth control compared with PBS and either single-agent treatment (**e**). Neither KROS-101 nor anti-PD-1 monotherapy significantly prolonged survival relative to PBS, whereas the combination significantly prolonged survival (**f**). Tumor growth was analyzed by mixed-effects modeling with Tukey’s multiple-comparisons test; survival was analyzed by Kaplan–Meier analysis with Šídák-adjusted pairwise log-rank (Mantel–Cox) tests. * p < 0.05; ** p < 0.01; *** p < 0.001; **** p < 0.0001. Data shown as mean ± s.e.m.

### GITRL stabilization reactivates antitumor immunity in patient-derived tumor explants (PDEs)

We finally asked whether pharmacological stabilization of endogenous GITRL within intact human tumor tissue is sufficient to reverse intratumoral immune suppression. To test this, freshly resected tumor specimens from five histological subtypes spanning primary brain tumors and brain metastases, including glioma, GBM, ependymoma, triple-negative breast cancer metastasis, and uterine carcinoma metastasis, were mechanically fragmented into PDEs and treated *ex vivo* with KROS-101 (Fig. 8a). KROS-101 increased the proportion of CD45⁺ cells among total viable cells and expanded CD8⁺ T cells within the CD45⁺ compartment, while concomitantly reducing FoxP3⁺ Tregs, resulting in a significantly elevated CD8⁺:Treg ratio across tumor subtypes (Fig. 8b-e and h). Within the CD8⁺ T cell compartment, KROS-101 increased the frequency of Tbet⁺ (Fig. 8f, k) and Tbet⁺TOX⁺ cells (Fig. 8f) with no significant reduction in TOX⁺ cells (Fig. 8l). It also expanded GZMB⁺Perforin⁺ cytotoxic effectors (Fig. 8g, j) and Ki67⁺ proliferating cells (Fig. 8i). KROS-101 treatment was associated with the emergence of a CD11c⁺XCR1⁺ cDC1-like population (Fig. 8m). Intratumoral cDCs exhibited upregulation of PD-L1 (Fig. 8n, p), CD86 (Fig. 8o, q), and HLA-DR (Fig. 8r). To determine whether these effects were reproducible within a single tumor histology, we performed an independent validation in additional PDEs restricted to GBM. KROS-101 produced a similar pattern of immune remodeling in this GBM-specific cohort, providing within-histology confirmation of the changes observed across the initial heterogeneous PDE cohort (Supplementary Fig. 7d-n). To test whether this activity required preservation of endogenous APC-T cell interactions, matched portions of the same tumors were enzymatically dissociated into single-cell suspensions, and we depleted CD14^+^/CD11c^+^ autologous APC fractions before treatment and performed functional assays across all five donors. APC depletion markedly reduced KROS-101-driven of cytotoxicity and the frequencies of IFN-γ and TNF-α expressing CD8^+^ cells in all specimens tested (Supplementary Fig. 7a-c). To determine whether GITRL stabilization alters Treg suppressive function itself, and whether this requires GITRL, we measured effector T cell proliferation in Teff:APC:Treg tricultures treated with KROS-101 in the presence or absence of a neutralizing anti-human GITRL antibody (Supplementary Fig. 7o-r). Tregs markedly suppressed Teff proliferation, and KROS-101 significantly relieved this suppression (Supplementary Fig. 7o). KROS-101 concurrently increased CD80+, CD86+ and MHC-II+ APCs (Supplementary Fig. 7p), reduced FoxP3+, IL-10+ and TGF-β+ Tregs (Supplementary Fig. 7q), and increased CD25+, TNF-α+, IFN-γ+, granzyme B+ and CD107a+ Teff cells (Supplementary Fig. 7r). Each of these effects, including the relief of Treg-mediated suppression, was abolished by GITRL neutralization, establishing that KROS-101 reduces Treg suppressive function through a GITRL-dependent mechanism.

**Figure 8:**
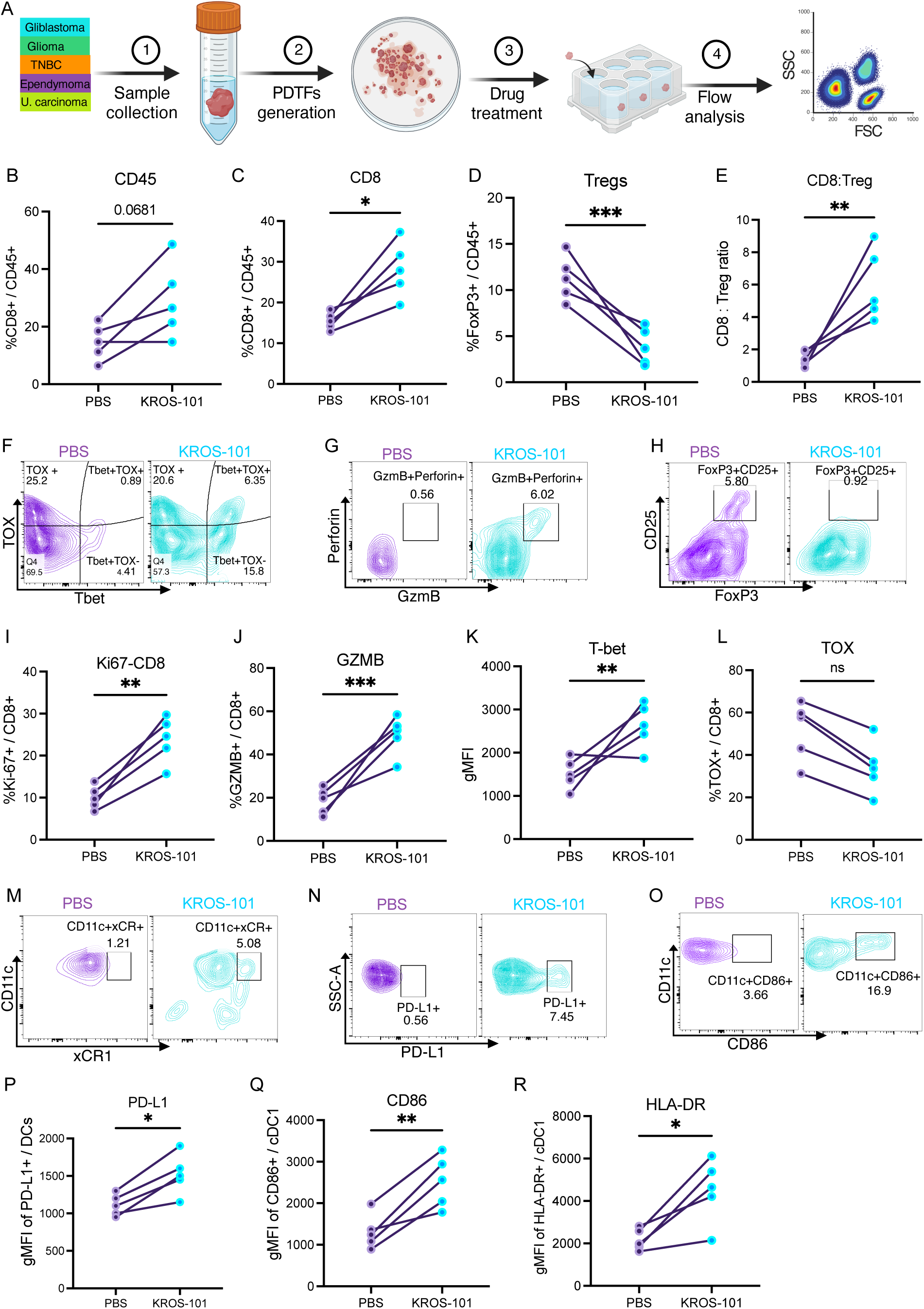
KROS-101 enhances effector CD8⁺ T cell activity and reduces Treg-mediated suppression in patient-derived tumor explants (PDEs). **a**, Workflow for analysis of PDEs from glioblastoma, glioma, ependymoma, triple-negative breast cancer (TNBC) brain metastasis, and uterine carcinoma brain metastasis. Tumor samples were processed to generate PDEs, treated with KROS-101 *ex vivo*, and analyzed by flow cytometry. **b-e**, Immune composition in PDEs following PBS or KROS-101 treatment. KROS-101 did not significantly alter the total CD45⁺ immune fraction (**b**) but increased the proportion of CD8⁺ T cells (**c**), reduced FoxP3⁺ Tregs (**d**), and increased the CD8:Treg ratio (**e**). **f-h**, Representative flow cytometry plots showing T-bet/TOX phenotypes (**f**), increased cytotoxic CD8⁺ T cells expressing GZMB and perforin (**g**), and decreased FoxP3⁺CD25⁺ Tregs (**h**) following KROS-101 treatment. **i-l**, Functional phenotype of CD8⁺ T cells in PDEs. KROS-101 increased proliferating Ki-67⁺ CD8⁺ T cells (**i**), enhanced GZMB expression (**j**), and increased T-bet expression (**k**), with no significant change in TOX expression (**l**). **m-o**, Representative flow cytometry plots showing the frequency of XCR1⁺CD11c⁺ cDC1 cells (**m**), PD-L1 expression (**n**), and CD86 expression on cDC1 cells (**o**). **p-r**, Quantification of DC activation markers showing increased PD-L1 expression on DCs (**p**), elevated CD86 expression (**q**), and increased HLA-DR expression on cDC1 cells (**r**) following KROS-101 treatment. Statistical comparisons were performed using paired t-test. ns, not significant; * p < 0.05; ** p < 0.01; *** p < 0.001. Each dot represents an individual patient PDE; lines connect matched PBS and KROS-101 conditions. n = 5 patient samples. The experimental design in panel A was created in BioRender. Admasu, T. (2026) https://BioRender.com/k1108md

## Discussion

The clinical failure of antibody-based GITR agonists (*6, 19, 20*) has been puzzling, given robust preclinical data. Our findings reveal a possible molecular explanation: antibodies bypass the trimeric architecture of GITRL, creating non-physiological receptor clustering that excludes APC-dependent costimulation and reverse signaling. Stabilizing endogenous trimeric or oligomeric GITRL addresses both deficiencies, thereby restoring the geometric constraints and cellular context required for productive GITR activation. This ligand-centric approach contrasts sharply with the receptor-targeting paradigm that has dominated TNF superfamily therapeutics. Multiple GITR antibodies, TRX518, MK-4166, AMG 228, and MEDI1873, advanced to clinical testing based on murine efficacy (*21–24*), yet all showed minimal activity in patients, even with checkpoint blockade (*8–10*). A critical oversight was the molecular divergence between human and mouse GITRL: human GITRL forms stable trimers, whereas mouse GITRL exists predominantly as dimers (*11*). This species difference fundamentally alters receptor engagement geometry and may explain why murine models overestimate antibody efficacy. Our use of humanized GITR/GITRL knock-in mice addresses this limitation and demonstrates that stabilizing the human trimeric ligand produces antitumor activity and immune remodeling that differ from antibody-mediated GITR agonism across B16-F10 and MC38. The orthotopic GL261 study extends these findings to an intracranial CNS tumor and demonstrates that KROS-101-mediated immune modulation can translate into a survival benefit in a native anatomical setting (Fig. 3h). Several features may distinguish ligand stabilization from antibody-mediated GITR agonism. First, the native GITR–GITRL interaction may promote receptor organization distinct from that induced by conventional bivalent antibodies, whose activity depends on geometry, valency, epitope, and, in some cases, Fc-mediated crosslinking (*12*). Second, ligand stabilization retains GITR signaling within the APC–T cell interface, alongside CD28–CD80/CD86 costimulation that supports T cell activation and effector differentiation (*25*). Finally, maintaining the native receptor– ligand interaction may permit bidirectional signaling through the GITR–GITRL axis. Although our functional analyses focused on CD8⁺ cytotoxic responses, the partial dependence of KROS-101-enhanced killing on MHC class II (Fig. 6f), together with the expression of GITR on both CD4⁺ and CD8⁺ T cells, raises the possibility that GITRL stabilization additionally engages antigen-experienced CD4⁺ T cells in an MHC-II-restricted manner. Neutralization of GITRL attenuated KROS-101-enhanced cytotoxicity, supporting a specific requirement for GITRL in its activity. This dependence may have implications for patient stratification, as responsiveness may be influenced by the availability of GITRL-expressing APCs within the tumor microenvironment. Consistent with this model, low-GITRL glioma and ependymoma TIL samples showed limited responsiveness, whereas baseline GITRL expression on macrophage and DC subsets correlated with KROS-101-enhanced cytotoxicity. Together, these findings support further evaluation of APC-associated GITRL as a potential biomarker of response to ligand stabilization.

The selective destabilization of intratumoral Tregs represents a mechanistically distinct outcome from antibody agonism. Although GITR is highly expressed on Tregs and its ligation was proposed to attenuate suppression (*7, 26*), antibody agonists minimally affected Treg function in patients (*8, 10*). In the autologous triculture system, KROS-101 relieved Treg-mediated suppression, and GITRL neutralization abolished this effect and the accompanying APC, Treg, and effector T cell phenotypic changes (Supplementary Fig. 7). Ligand stabilization creates an inflammatory niche enriched in TNFα, IFN-γ, and APC-derived costimulation that likely exceeds the threshold for Treg transcriptional destabilization. This is consistent with recent evidence that Treg identity is fragile under strong inflammatory pressure (*27, 28*) and that IFN-γ disrupts Treg suppressive programs (*27*). Importantly, Treg depletion occurred selectively within tumors, not systemically, suggesting tumor-specific microenvironmental factors, hypoxia, adenosine, or metabolic stress, cooperate with GITR signaling to preferentially destabilize intratumoral Treg fitness. KROS-101-treated tumors exhibited organized APC-T cell clusters, hallmarks of lymphoid neogenesis (*29, 30*). This raises the possibility that ligand stabilization converts immune-excluded tumors into organized lymphoid microenvironments permissive to adaptive immunity. The ligand-stabilizing paradigm established here may extend to other TNF receptor superfamily costimulatory molecules. Agonist antibodies targeting 4-1BB (CD137), OX40 (CD134), and CD27 have encountered analogous clinical challenges despite promising preclinical data (*31–33*). Each receptor engages trimeric ligands expressed on APCs and exhibits ligand geometry-dependent signaling (*34*). Stabilizing 4-1BBL, OX40L, or CD70 could restore physiological receptor clustering, preserve APC-dependent costimulation, and enable reverse signaling-mediated APC activation, offering a generalizable strategy for salvaging costimulatory pathways that have proven refractory to antibody-based targeting. Because KROS-101 selectively stabilizes human trimeric GITRL and does not engage murine dimeric GITRL, all *in vivo* studies were conducted in hGITR/hGITRL humanized knock-in mice. Most efficacy models were subcutaneous, but the orthotopic GL261 study provides initial validation in a CNS setting; additional orthotopic models remain warranted. Notably, the patient-derived tumor explant experiments (Fig. 8) interrogate KROS-101 within intact, architecturally preserved human tumor tissue and thus provide a complementary, translationally relevant readout. From a translational and manufacturing standpoint, the small-molecule nature of KROS-101 offers scalability and cost advantages over biologic agonists, and the persistence of CD8⁺ T cell reprogramming after drug removal (Fig. 6g-m) indicates that durable antitumor benefit need not require continuous target engagement. In summary, we establish ligand stabilization as a mechanistically distinct approach to therapeutic GITR agonism that overcomes molecular and cellular limitations inherent to antibody-mediated receptor crosslinking. By preserving GITRL-dependent APC-T cell interactions, the ligand-stabilizing paradigm coordinates APC maturation with T cell activation and reduces Treg-mediated suppression to support antitumor immunity. These findings provide a conceptual framework for targeting TNF receptor superfamily costimulatory pathways by modulating their native ligands and establish a mechanistic rationale for advancing ligand-stabilizing pharmacology into clinical development for treatment-refractory solid tumors.

## Acknowledgements

The authors thank the patients and healthy volunteers who generously donated samples for this research; without their contributions, this work would not have been possible. We thank the surgical and clinical teams at Cedars-Sinai Medical Center for their assistance with tumor specimen collection and patient consent coordination. We acknowledge the developers and maintainers of publicly available datasets from TCGA and TISCH2, which enabled the bioinformatics analyses in this study.

## Author contributions

T.D.A. designed and supervised all experimental work, performed in vitro KROS-101 characterization, tri-culture and APC-T cell co-culture experiments, TIL isolation and characterization, patient-derived tumor explant studies, flow cytometry analyses, imaging mass cytometry experiments, cell segmentation, and neighborhood enrichment quantification, and drafted the manuscript. K.S. and S.Y. performed in vitro cell-based assays. D.K., H.W. and K.W. involved in *in vivo* tumor experiments in hGITR/hGITRL knock-in mice, including tumor implantation, treatment administration, tumor measurement, and survival monitoring. M.K conducted surface plasmon resonance binding studies. J.R. contributed to human tumor specimen acquisition and clinical coordination. M.R. provided scientific direction, oversaw the drug design aspects of the study, and contributed to manuscript revision. J.Y. supervised the overall project, provided strategic direction, secured funding, and revised the manuscript. All authors reviewed and approved the final manuscript.

## Data, Code and Material Availability

All data needed to evaluate and reproduce the results in the paper are present in the paper and/or the Supplementary Materials. This study did not generate new code. KROS-101 can be provided by Kairos Pharma, Ltd pending scientific review and a completed material transfer agreement with Kairos Pharma, Ltd. Requests for KROS-101 should be submitted to: Ramachandran Murali at.

## Methods and Materials

***Sex as a biological variable.*** Both male and female mice were included, and data were pooled; sex was not considered as a biological variable for animal models. Human samples included both sexes where available, and sex-stratified analyses were not performed.

### TCGA and single-cell transcriptomic analysis

Bulk tumor and matched normal tissue expression data for *TNFSF18* and *TNFRSF18* were retrieved from The Cancer Genome Atlas (TCGA) PanCancer Atlas using the TCGA online portal (https://www.cancer.gov/tcga). Tumor ImmuneScore and StromalScore were computed from TCGA RNA-seq data using the ESTIMATE web tool (https://bioinformatics.mdanderson.org/estimate/). Gene set enrichment analysis (GSEA) was performed using the MSigDB web interface (https://www.gsea-msigdb.org/gsea/msigdb/) against Hallmark and IFN-γ response gene sets. Single-cell RNA-seq data from publicly available pan-cancer tumor atlases were accessed and queried through the CancellDB (https://cancelldb.org) and Tumor Immune Single-cell Hub (TISCH2; http://tisch.comp-genomics.org) web portals. Cell type annotations were adopted from the original publications as curated within each portal. *TNFSF18* expression across annotated cell clusters was visualized as dot plots and feature plots using the built-in visualization tools of the GEPIA2 and TIMER2 portals. Kaplan–Meier survival analysis was performed on TCGA lower-grade glioma (LGG) patients stratified by high/low expression of *TNFSF18* and *TNFRSF18*, or both, using the KM-plot web tool (https://kmplot.com), with the optimal cutpoint determined automatically by the platform. Log-rank tests were used to assess significance, with false discovery rate correction applied for multiple comparisons.

### KROS-101: structure-based design and compound identification

Atomic structures of hGITRL were obtained from Protein Data Bank (PDB: 2Q1M and 3B94). KROS-101 was identified as a small-molecule GITR/GITRL modulator through a structure-guided design strategy using the Glide protocol (Schrodinger, Inc) using default parameters as described previously (*35*). The design of GITR/GITRL modulator compounds was based on the crystal structure of the GITR/GITRL complex, which enabled identification of the GITRL binding interface as a tractable target for small-molecule intervention. Small molecules that stabilize the GITR-GITRL receptor complex were identified iteratively through biophysical screening assays. KROS-101 was selected from this program based on its selectivity for GITRL and its ability to modulate the GITR/GITRL immune regulatory axis. KROS-101 has a chemical formula of C_22_H_28_N_2_O_6_ and a molecular weight of 416 g/mol.

Its 2D structure is shown in Supplementary Fig. 2f and the IUPAC Chemical Name is : 5,5’-([1,1’-biphenyl]-4,4’-diylbis(azanediyl))bis(2 (hydroxymethyl)tetrahydrofuran-3-ol) (Product ID: 5483007; SMILES: OC1CC(OC1CO)NC2=CC=C(C=C2)C(C=C3)=CC=C3NC(CC4O)OC4CO) It was identified from the ChemBridge (San Diego, CA) chemical screening library and synthesized by ChemBridge Corporation as a racemic mixture at >95% purity, as confirmed by vendor quality control analysis. For in vitro experiments, KROS-101 was prepared as a 6 mM stock solution in DMSO and further diluted in PBS to achieve the indicated working concentrations, with DMSO maintained at ≤0.1% (v/v) in all biological assays. Aqueous solubility at working concentrations was confirmed prior to use.

### GITRL binding affinity: Surface Plasmon Resonance (SPR)

The GITRL binding affinity of KROS-101 was determined using surface plasmon resonance using a Sartorius Octat SF3 instrument. Briefly, recombinant hGITRL was immobilized on a HisCap sensor chip to 22,000 RU. hGITRL small molecules (KROS-series) were analyzed at serial doubling concentrations maximized at 100 μM in PBST buffer containing 5% DMSO (pH 7.4). Analytes were injected at 10 μL/min for 3 min at room temperature. The final results were obtained using analysis software provided by the vendor. For the dimeric-ligand comparator experiment, recombinant mouse GITRL-Fc (Novus Biologicals, catalog no. NBP2-26579) was immobilized on the sensor surface, and KROS-101 binding was evaluated at two concentrations using the same running buffer, acquisition, reference-subtraction, and analysis procedures applied to human GITRL.

### NF-κB luciferase reporter assay

NF-κB signaling was measured using Jurkat NF-κB luciferase reporter cells. Cells were seeded at approximately 4 x 10^^4^ cells per well in white 96-well plates and stimulated with recombinant trimeric human GITRL in the presence or absence of KROS-101. For the ligand-limited experiment shown in Fig. 1d, GITRL was used at 125 ng/mL; 1 µg/mL GITRL was used as the saturating-ligand condition. Following 4 h of stimulation at 37°C, firefly luciferase activity was measured using a luciferase detection reagent and a luminescence plate reader. Background luminescence from cell-free wells was subtracted, and NF-κB activity was expressed relative to the vehicle-treated control. Experiments were performed in three independent replicates (n = 3). Statistical comparisons were performed using one-way ANOVA with Tukey’s multiple-comparisons test.

### Human PBMC isolation and primary cell purification

Peripheral blood mononuclear cells (PBMCs) were isolated from healthy donors by density-gradient centrifugation over Ficoll-Paque PLUS (GE Healthcare). Human CD8⁺ T cells, CD4⁺CD25⁺FoxP3⁺ regulatory T cells (Tregs), monocyte-derived dendritic cells (moDCs), and monocyte-derived M1 macrophages were purified from autologous PBMCs of the same donor for each experiment. CD8⁺ T cells were enriched by negative magnetic selection (Miltenyi Biotec). Tregs were enriched using the CD4⁺CD25⁺ Regulatory T Cell Isolation Kit (Miltenyi Biotec). Monocytes were isolated by CD14⁺ positive selection (Miltenyi Biotec) and differentiated into moDCs by culture with IL-4 (50 ng/mL) and GM-CSF (100 ng/mL) for 6 days in RPMI-1640 with 10% heat-inactivated FBS, or into M1 macrophages by culture with M-CSF (50 ng/mL) for 5 days followed by 24 h polarization with IFN-γ (20 ng/mL) and LPS (100 ng/mL). All cultures were maintained at 37°C in 5% CO₂. Cell purity was confirmed by flow cytometry before each experiment.

### Tri-culture system and GITRL surface stabilization assay

Tri-culture systems were established in 96-well plates at a ratio of 5:1:2 (CD8⁺ T cells:Tregs:APCs) in complete RPMI-1640. KROS-101 or DMSO vehicle was added at the indicated concentrations at the time of co-culture initiation. For APC monoculture conditions, moDCs or M1 macrophages were cultured alone with KROS-101 or vehicle and sampled at the indicated time points. To assess GITRL surface stabilization, cells were stained with anti-human GITRL antibody at the indicated time points (0, 6, 12, 24, 48, and 72 h) without prior fixation, on ice, to detect surface-resident protein. GITRL geometric mean fluorescence intensity (gMFI) was measured by flow cytometry and normalized to vehicle-treated controls.

### APC-T cell co-culture and bidirectional activation assay

For bidirectional activation experiments, autologous moDCs or M1 macrophages were co-cultured with purified CD8⁺ T cells and Tregs (as above) in the presence of KROS-101 or DMSO vehicle for 48 h. Cells were harvested and processed for intracellular cytokine staining: stimulated for 4 h with PMA (50 ng/mL) and ionomycin (500 ng/mL) in the presence of Brefeldin A (5 µg/mL), then surface-stained, fixed, and permeabilized (eBioscience FoxP3/Transcription Factor Staining Buffer Set) for intracellular detection of IFN-γ, TNF-α, IL-10, TGF-β, and FoxP3. For the APC-exclusion condition, Tregs and CD8⁺ T cells were co-cultured in the absence of any APC population to confirm APC requirement for KROS-101 activity. For functional suppression assays, effector T cell proliferation was quantified in matched Teff:APC and Teff:APC:Treg cultures treated with vehicle or KROS-101, with or without GITRL neutralization.

### GITRL neutralization blockade

To confirm GITRL dependence, autologous tumor-killing and Teff:APC:Treg suppression assays were conducted in the presence of a GITRL-neutralizing antibody or isotype control. The neutralizing antibody was added 30 min before KROS-101 treatment. Effector T cell proliferation, APC activation, Treg phenotype, effector T cell activation and cytotoxicity markers, cytokine production, and tumor cell killing were assessed as described above and below.

### TIL isolation and profiling from patient tumor specimens

Freshly resected tumor specimens from patients with glioma, glioblastoma (GBM), triple-negative breast cancer, ependymoma, or uterine carcinoma were obtained following informed consent under IRB protocol Pro00008922. Specimens were processed within 2 h of surgical resection. Tissue was mechanically dissociated through 70 µm cell strainers in complete RPMI-1640, and tumor-infiltrating lymphocytes (TILs) were enriched by density-gradient centrifugation over Ficoll-Paque. For GITRL expression profiling, freshly isolated TILs were stained with a multicolor flow cytometry panel targeting immune subset markers and GITRL (full panel provided in Supplementary Table 2). DC subsets were identified as Lin⁻ (CD3⁻CD19⁻CD56⁻) HLA-DR⁺CD11c⁺ cells; M1-like macrophages as CD14⁺CD68⁺CD80⁺/CD86⁺ cells. GITRL expression was quantified as gMFI on each APC subset and correlated with subsequent functional readouts.

### Ex vivo autologous tumor killing assay (patient TILs)

For autologous killing assays, freshly isolated patient TILs were co-cultured with autologous primary tumor cells at a 10:1 effector-to-target (E:T) ratio in 96-well flat-bottom plates in the presence of KROS-101 or vehicle control. Autologous tumor cells were plated 24 h before co-culture to allow adherence. Immediately following TIL addition, plates were transferred to an Incucyte S3 live-cell analysis system (Sartorius) and imaged every 2–3 h for the duration of the assay. Tumor cell killing was quantified by longitudinal measurement of incorporation of a cell-impermeant dead-cell reagent (Incucyte Cytotox Red). Cytotoxicity was calculated as the reduction in viable target cell area relative to vehicle-treated wells and normalized to tumor-only controls. Killing kinetics were analyzed as percent target cell loss over time and area-under-the-curve (AUC) relative to tumor-only wells.

### Animal models

Humanized double knock-in mice co-expressing human GITR and human GITRL ectodomains, referred to as hGITR/hGITRL mice, were obtained from GenOway (Lyon, France). The hGITR/hGITRL knock-in line was generated on a C57BL/6J background using C57BL/6 mice as the parental strain. In this model, the extracellular domains of endogenous murine GITR and GITRL are replaced by their human counterparts while preserving intact murine transmembrane and intracellular signaling domains and native promoter control. Mice were housed under specific pathogen-free conditions with ad libitum access to food and water under a 12-h light/dark cycle. Age-matched, mixed-gender mice (6-8 weeks) were used for all tumor experiments. All procedures were approved by the Cedars-Sinai Medical Center Institutional Animal Care and Use Committee (IACUC protocol IACUC008158) and conducted in accordance with the NIH Guide for the Care and Use of Laboratory Animals. All procedures, including tumor implantation, drug administration, and euthanasia, were performed in accordance with approved protocols and institutional guidelines. hGITR/hGITRL expression across immune populations was validated by flow cytometry on peripheral blood and splenocytes before tumor experiments.

### Tumor cell lines and implantation

MC38 (murine colon adenocarcinoma), B16-F10-Luc2 (luciferase-expressing murine melanoma), and YUMM1.7 (murine melanoma) cells were obtained from ATCC and maintained in DMEM supplemented with 10% FBS and 1% penicillin-streptomycin at 37°C in 5% CO₂. Cells were verified as mycoplasma-negative by PCR and passaged fewer than 10 times after receipt. For MC38 and YUMM1.7 experiments, 1×10⁵ cells in 100 µL PBS were injected subcutaneously into the right flank. All tumor efficacy experiments used group sizes of 8-10 mice per treatment arm, determined by a priori power analysis (80% power, two-sided α = 0.05) using pilot tumor growth data from our laboratory. KROS-101 or vehicle treatment was initiated 3 days post-implantation. For B16-F10-Luc2 experiments, 1×10⁵ cells in 100 µL PBS were injected subcutaneously, and KROS-101 or vehicle treatment was initiated 2 days post-implantation. Tumor volumes were measured with a digital caliper every 2-3 days and calculated as (length × width²)/2. Bioluminescence imaging was performed using the IVIS Spectrum system (PerkinElmer) following intraperitoneal injection of D-luciferin (150 mg/kg), with signal quantified as total flux (photons/s) in the tumor region using Living Image software. Mice were euthanized when tumor volume reached 2,000 mm³ or upon development of ulceration or clinical distress.

**GL261 intracranial tumor implantation:** Anesthetized mice were secured in a stereotactic frame, and GL261 cells (4 × 10⁴ cells in 4 μl PBS) were implanted intracranially into the right cerebral hemisphere at 2 mm lateral to the midline, 2 mm anterior to the coronal suture, and a depth of 3.5 mm. The needle was retained for 5 min before withdrawal. For survival studies, mice were monitored according to protocol-defined neurologic and clinical humane endpoints.

### KROS-101 treatment administration

KROS-101 was dissolved in DMSO and diluted in PBS to yield a 6 mg/mL stock immediately before administration. Mice received KROS-101 20 mg/kg or vehicle by intraperitoneal injection every other day for a total of seven doses, beginning on the indicated treatment day. Group allocation was performed by simple randomization; investigators were not blinded to treatment allocation during tumor measurements.

**Checkpoint-blockade combination studies:** For checkpoint-blockade combination studies, hGITR/hGITRL double-knock-in mice bearing MC38 or B16-F10 tumors were monitored using the tumor-measurement and humane-endpoint procedures described above. Mice received PBS, KROS-101, checkpoint antibody, or the corresponding combination. Anti-PD-1 (clone RMP1-14, Bio X Cell; 200 μg per mouse, intraperitoneally) was evaluated in the MC38 and B16-F10 models, and anti-CTLA-4 (clone 9H10, Bio X Cell; 100 μg per mouse, intraperitoneally) was evaluated in the MC38 model. KROS-101 dosing followed the model-specific in vivo regimen described above. Checkpoint antibodies were administered intraperitoneally, with the same antibody schedule used in the corresponding monotherapy and combination groups. Tumor growth and survival were analyzed as described under Statistical analysis.

### Flow cytometry

Tumors were excised and mechanically dissociated through 70 µm cell strainers in digestion buffer (RPMI containing collagenase IV [1 mg/mL; Sigma] and DNase I [0.1 mg/mL; Sigma]) for 30 min at 37°C with agitation. Single-cell suspensions were filtered through 40 µm strainers, washed, and subjected to red blood cell lysis (ACK buffer, 2 min, RT). Spleens were mechanically dissociated through 70 µm strainers and similarly processed. Peripheral blood was collected by retro-orbital bleeding into EDTA-coated tubes; red blood cells were lysed before staining. Single-cell suspensions were incubated with TruStain FcX (anti-mouse CD16/32; BioLegend) for 10 min on ice, followed by surface staining in FACS buffer (PBS + 2% BSA + 0.05% sodium azide) for 20 min on ice. For intracellular cytokine staining, cells were restimulated ex vivo with PMA (50 ng/mL) and ionomycin (500 ng/mL) with Brefeldin A (5 µg/mL) for 4 h at 37°C prior to staining. Fixation and permeabilization were performed using the eBioscience FoxP3/Transcription Factor Staining Buffer Set per the manufacturer’s protocol. Samples were acquired on a Sony ID7000 spectral flow cytometer. Spectral unmixing was performed using single-color controls and the ID7000 Software Suite. Data analysis was performed using FlowJo v10.9 (BD Biosciences). Immune populations were identified using the hierarchical gating strategy provided in Supplementary Fig. 3. Full antibody panel details are provided in Supplementary Table 1.

### Ex vivo cytotoxicity assay

CD45⁺ TILs were enriched from digested tumors using the EasySep Mouse CD45 Positive Selection Kit (STEMCELL Technologies), followed by CD8⁺ T-cell enrichment using the EasySep Mouse CD8 Positive Selection Kit. MC38 or B16-F10-Luc2 target cells (1 × 10³ cells per well) were seeded in 96-well plates and allowed to adhere for 4 h. CD8⁺ TILs were added at an effector-to-target ratio of 10:1 in complete RPMI-1640. Tumor cell killing was monitored for 24-48 h using an Incucyte SX5 system (Sartorius) with a caspase-3/7 apoptosis reagent and quantified by caspase-3/7 fluorescence. For mechanistic studies, anti-IFN-γ (clone XMG1.2; 10 µg/ml), concanamycin A (100 nM), anti-MHC class I (clone M1/42; 20 µg/ml), or anti-MHC class II (clone M5/114.15.2; 20 µg/ml) was added at the start of coculture. Isotype-matched antibodies were used as controls.

### Antigen-specific splenocyte recall response

Spleens were harvested from vehicle-or KROS-101-treated tumor-bearing mice at day 10 post-treatment initiation. Single-cell suspensions were prepared and plated at 2×10⁵ cells per well in 96-well plates. APCs pulsed with MC38 cell lysate were added as restimulation antigen. For MHC-I blockade experiments, anti-MHC class I antibody (clone M1/42; 20 µg/mL) or isotype control was added at the start of restimulation. After 12 h, Brefeldin A (5 µg/mL) was added for an additional 4 h. IFN-γ production by CD8⁺ T cells was measured by intracellular cytokine staining and flow cytometry.

### Durable reprogramming co-culture assay

To assess whether KROS-101-induced functional changes persist after drug removal, CD8⁺ TILs and Tregs were isolated separately from tumors of PBS-or KROS-101-treated mice on day 10. CD8⁺ T cells were sorted by EasySep Mouse CD8 Positive Selection Kit (STEMCELL Technologies). Tregs were enriched by CD4⁺CD25⁺ positive selection. Cells from each treatment group were combined in four cross-over conditions: KROS-101-CD8⁺ alone, KROS-101-CD8⁺ + PBS-Tregs, PBS-CD8⁺ alone, and PBS-CD8⁺ + PBS-Tregs, at a 5:1 CD8:Treg ratio. Co-cultures were restimulated with anti-CD3 (1 µg/mL) and anti-CD28 (2 µg/mL) for 72 h. CD8⁺ cytotoxicity against MC38 or B16-F10 target cells was assessed at E:T = 10:1 using the Incucyte system. Intracellular cytokine accumulation of IL-2, TNF-α, IFN-γ and exhaustion marker expression (PD-1, LAG-3, TIM-3) was assessed by flow cytometry on harvested CD8⁺ T cells.

### Imaging mass cytometry (IMC)

B16-F10-Luc2 tumors were harvested from vehicle-or KROS-101-treated hGITR/hGITRL mice on day 7 post-treatment initiation. Tumors were fixed in 10% neutral-buffered formalin for 24 h at room temperature, processed, and embedded in paraffin (FFPE). Sections of 4 µm thickness were cut, deparaffinized, and subjected to heat-induced epitope retrieval in Tris-EDTA buffer (pH 9.0) at 95°C for 30 min. Tissue sections were blocked with 3% BSA/PBS for 1 h at room temperature. Antibodies were applied by incubating sections overnight at 4°C with all metal-conjugated antibodies simultaneously. Antibodies were conjugated to isotopically pure lanthanide metals using the Maxpar X8 Multimetal Labeling Kit (Standard BioTools) per the manufacturer’s protocol. After staining, sections were washed and stained with Iridium intercalator (250 µM; Standard BioTools) for 30 min to label nuclei. Sections were air-dried and ablated on a Hyperion Imaging System (Standard BioTools) at a laser frequency of 200 Hz with 1 µm resolution. Regions of interest (minimum 500×500 µm) were selected to capture the tumor core and infiltrating margin.

### Patient-derived tumor explant (PDE) preparation and ex vivo treatment

Fresh tumor tissue was obtained from patients undergoing surgical resection under IRB protocol Pro00008922 with written informed consent. Tissue was transported in cold RPMI-1640 and processed immediately. Samples were mechanically sectioned into fragments of ∼1 mm³ using sterile scalpels on ice. Fragments were distributed into 96-well round-bottom plates (7-8 fragments per well) in complete RPMI-1640 (RPMI + 10% FBS + 1% penicillin-streptomycin + 50 IU/mL IL-2) and treated immediately with KROS-101 or vehicle for 72 h at 37°C in 5% CO₂. At 72 h, explant fragments were dissociated by gentle vortexing in enzyme-free dissociation buffer (STEMCELL Technologies) followed by passage through 70 µm strainers. Single-cell suspensions were processed for flow cytometry as described above. For APC-depletion experiments, matched portions of the same tumor specimens were enzymatically dissociated (as in TIL isolation), and APCs were depleted using anti-human HLA-DR MicroBeads (Miltenyi Biotec) followed by LD column depletion. APC-depleted cell suspensions were then treated with KROS-101 or vehicle in suspension culture, and CD8⁺ T cell activation was assessed by flow cytometry.

**GBM-specific patient-derived explant validation:** An independent patient-derived explant cohort restricted to glioblastoma was processed, treated with KROS-101, and analyzed using the same explant-culture and flow-cytometry procedures described above. Analyses were performed at the donor level, with each tumor specimen treated as an independent biological replicate.

### Flow cytometric analysis of GITR and GITRL surface binding

To assess surface GITR and GITRL expression and their modulation across conditions, cells from tri-culture or TIL experiments were harvested, washed in cold FACS buffer, and incubated with human FcR Blocking Reagent (Miltenyi Biotec) for 10 min on ice. Cells were stained with directly conjugated anti-human GITR and GITRL antibodies for 30 min on ice. Viable cells were identified using a fixable viability dye (eBioscience Fixable Viability Dye eFluor 780). Surface markers, including CD3, CD4, CD8, CD14, CD11c, and HLA-DR, were included to identify cell populations. GITR and GITRL surface expression was quantified as gMFI.

### Quantification and statistical analysis

All statistical analyses were performed in GraphPad Prism (v10.0). For comparisons between two groups, the unpaired two-tailed Student’s t-test was used for normally distributed data (assessed by the Shapiro-Wilk test) and the Mann-Whitney U-test for non-normally distributed data. For comparisons among three or more groups, one-way ANOVA with Tukey’s post-hoc test (parametric) or Kruskal-Wallis with Dunn’s post-hoc test (non-parametric) was applied. For *in vivo* tumor growth curves, two-way ANOVA with Bonferroni correction was used. Kaplan-Meier survival curves were compared by log-rank (Mantel-Cox) test. Spearman’s rank correlation was used for association analyses between GITRL expression and cytotoxicity. All data are presented as mean ± SEM unless otherwise stated. Sample sizes (n) refer to biological replicates (individual mice or human donors) unless specified otherwise; exact n values for each panel are reported in figure legends. Statistical significance thresholds: *p < 0.05, **p < 0.01, ***p < 0.001, ****p < 0.0001; ns, not significant.

## Supplementary materials

**Supplementary Fig. 1:**
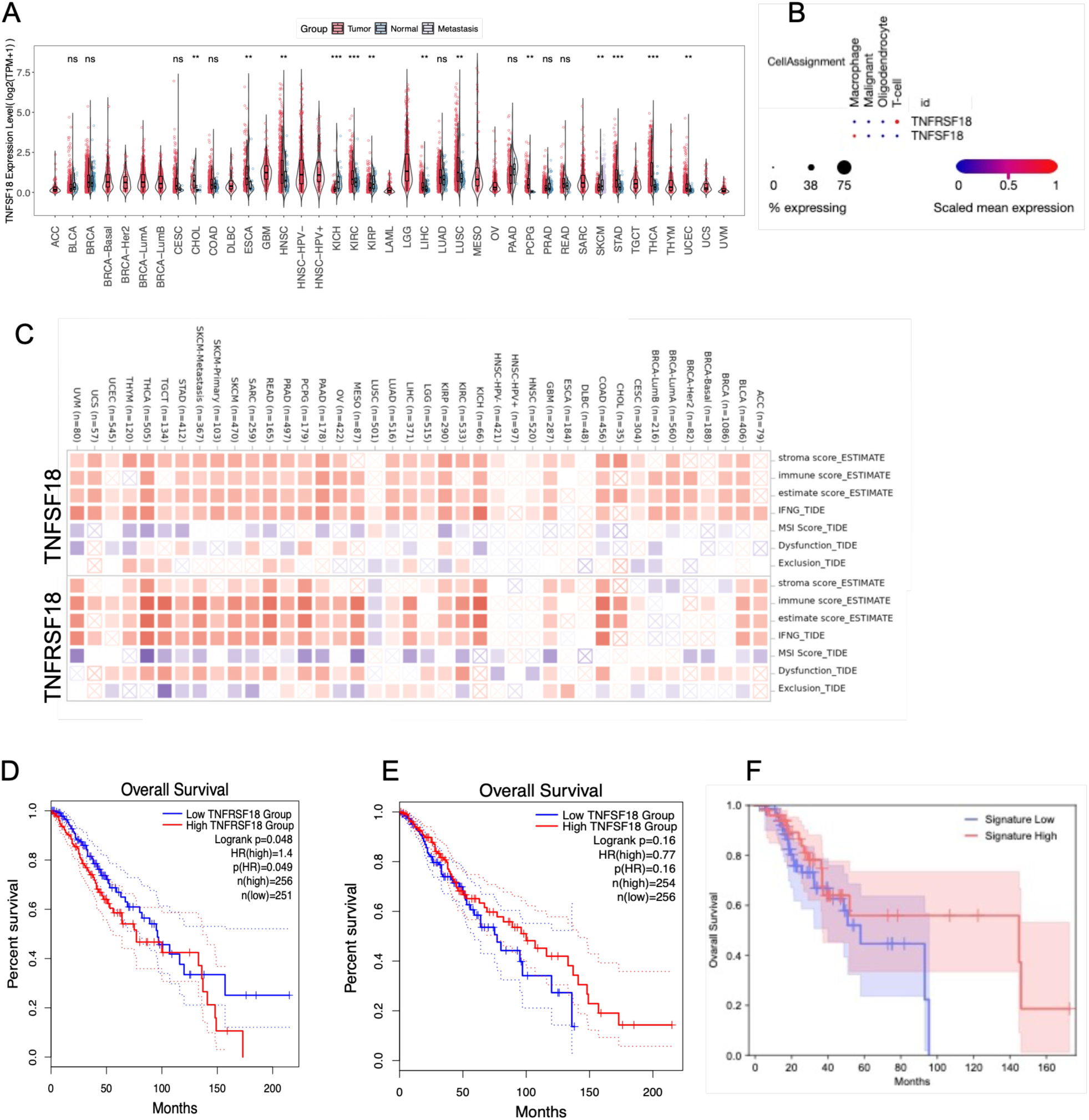
Pan-cancer expression and clinical relevance of the GITR-GITRL axis. **a**, Pan-cancer analysis of *TNFSF18* (GITRL) expression across tumor types from TCGA datasets showing relative expression levels in tumor, normal, and metastatic tissues. **b**, Cell-type specific expression of *TNFRSF18* (GITR) and *TNFSF18* (GITRL) across immune populations, highlighting enrichment in myeloid and lymphoid compartments. Dot size indicates the percentage of cells expressing the gene and color represents scaled mean expression. **c**, Correlation of TNFRSF18 and TNFSF18 expression with immune microenvironment features across cancers, including stromal score, immune score, interferon-γ signature, microsatellite instability (MSI), TIDE dysfunction score, and T cell exclusion metrics. **d**, Kaplan-Meier overall survival analysis stratified by TNFRSF18 expression levels in TCGA patient cohorts. **e**, Kaplan-Meier overall survival analysis stratified by TNFSF18 expression levels. **f**, Survival analysis based on a combined GITR-GITRL gene signature, comparing patients with high or low signature scores.

**Supplementary Fig. 2:**
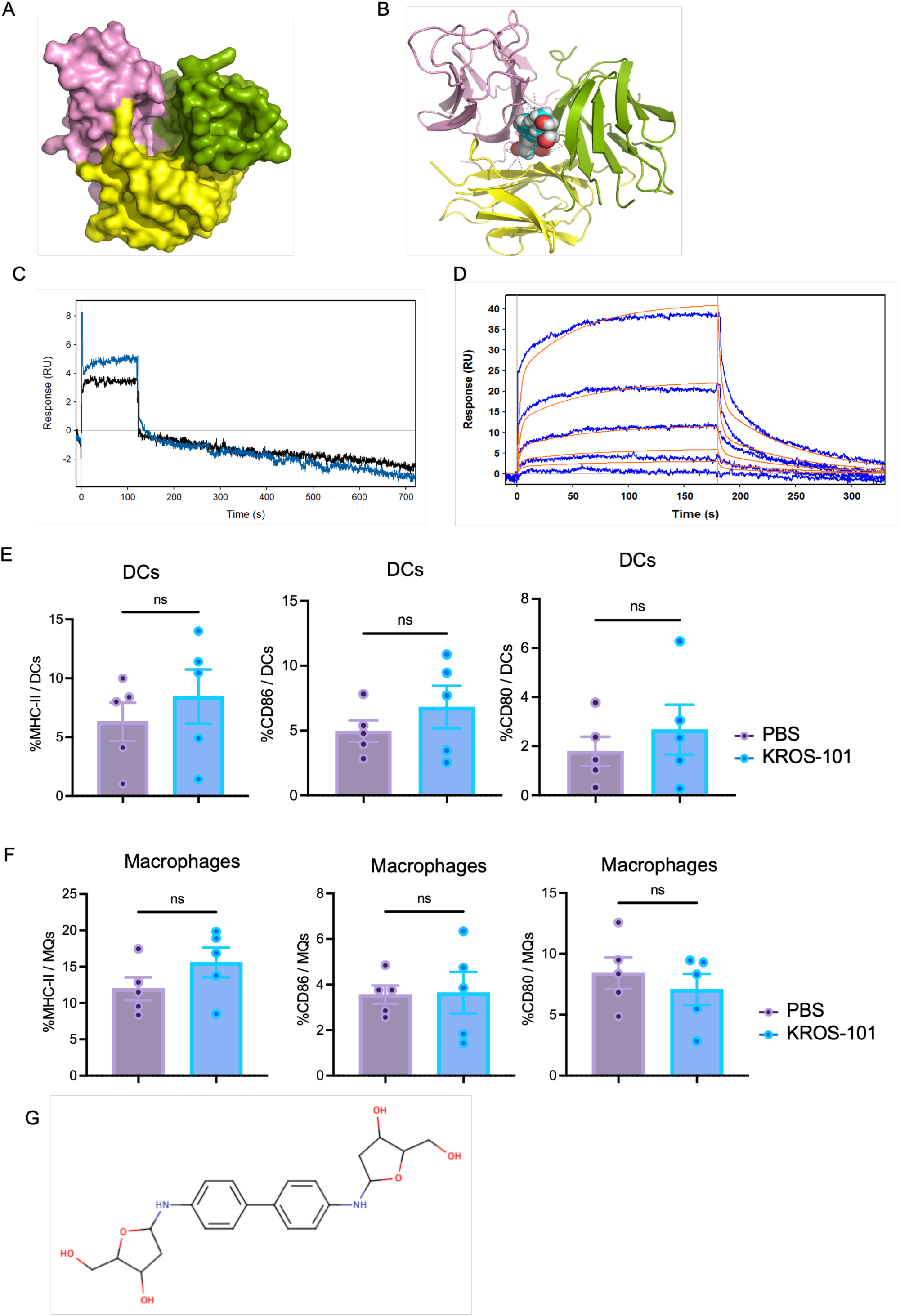
KROS-101 does not directly alter baseline APC activation markers. **a**, **Molecular Structure of human GITRL**: Atomic structure of human GITRL (PDB: 2Q1M and 3B94) shown as a molecular surface model. The three GITRL protomers are shown in pink, green, and yellow, highlighting the unique central cavity formed by the trimeric assembly. **b, Molecular model of KROS-101 bound to the trimeric GITRL**. Molecular docking of KROS-101 into the trimeric hGITRL interface. KROS-101 is shown as a space-filling model, with atoms colored by type (oxygen, red; nitrogen, blue; carbon, cyan; hydrogen, white). The model was generated using PyMOL (Morin et al., eLife, 2013). **c**, SPR analysis of KROS-101 binding to dimeric mGITRL-Fc. KROS-101 binding to immobilized mGITRL-Fc was evaluated by surface plasmon resonance (SPR) at two concentrations. KROS-101 produced a minimal binding response with rapid dissociation. **d, Direct binding of KROS-101 to hGITRL**. KROS-101 binding to hGITRL is evaluated by SPR. Dose-dependent binding of KROS-101 is shown in the blue sensogram curves. Langmuir model simulation (1:1) used to derive kinetic parameters of the bimolecular interactions shown in red. **e**, Frequency of DCs expressing MHCII, CD86, and CD80 following treatment with PBS or KROS-101. No significant differences were observed, indicating that KROS-101 does not directly induce classical DCs maturation. **f**, Expression of antigen-presentation and co-stimulatory markers in macrophages following treatment with PBS or KROS-101, including MHCII, CD86 and CD80. No significant changes were detected. Data represent flow cytometric analysis of antigen-presenting cells following treatment. Statistical comparisons were performed using two-tailed tests. **g**, 2D structure of KROS-101

**Supplementary Fig. 3:**
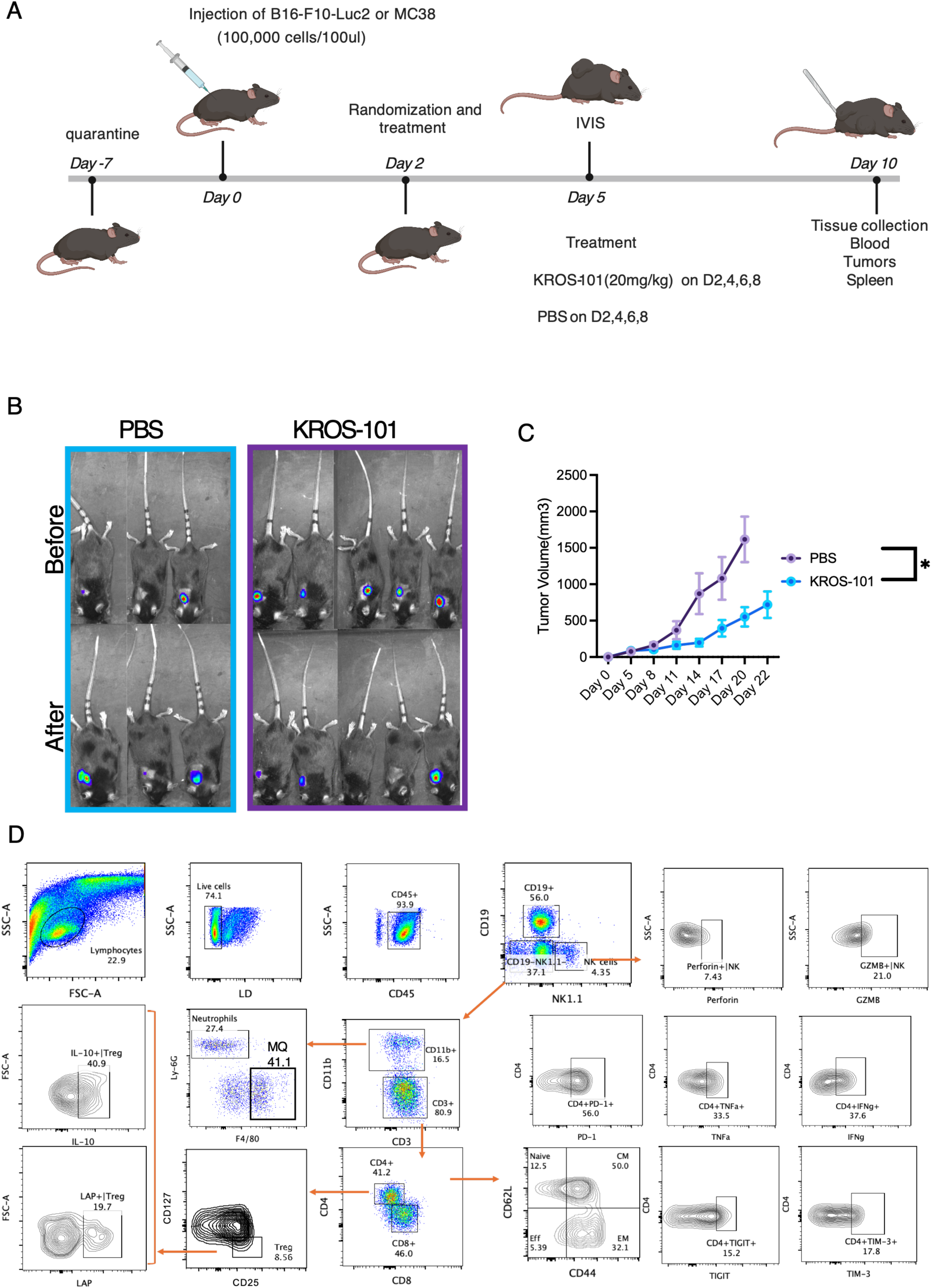
Experimental design and immune profiling of tumor-bearing mice treated with KROS-101. **a**, Schematic of the *in vivo* experimental workflow. Mice were injected with B16-F10-Luc2 or MC38 tumor cells (100,000 cells in 100 μl), followed by IVIS imaging. Mice were randomized and treated with KROS-101 (20 mg kg⁻¹) or PBS on days 7, 9, 11 and 13. Tumors, blood and spleens were collected for downstream analyses. **b**, Representative IVIS bioluminescence images of B16-F10-Luc2 tumor-bearing mice (used for flow analysis) before and after treatment with PBS or KROS-101, demonstrating reduced tumor signal in KROS-101-treated mice. **c**, Tumor growth curves showing reduced tumor volume in KROS-101–treated mice compared with PBS controls over the course of treatment. **d**, Flow cytometry gating strategy used for immune profiling of tumor-infiltrating leukocytes. Sequential gating identifies major immune populations, including lymphocytes, CD45⁺ leukocytes, NK cells, macrophages, neutrophils, CD3⁺ T cells, CD4⁺ and CD8⁺ T cells, regulatory T cells, and functional markers including perforin, granzyme B, PD-1, TNF-α, IFN-γ, TIGIT and TIM-3.

**Supplementary Fig. 4:**
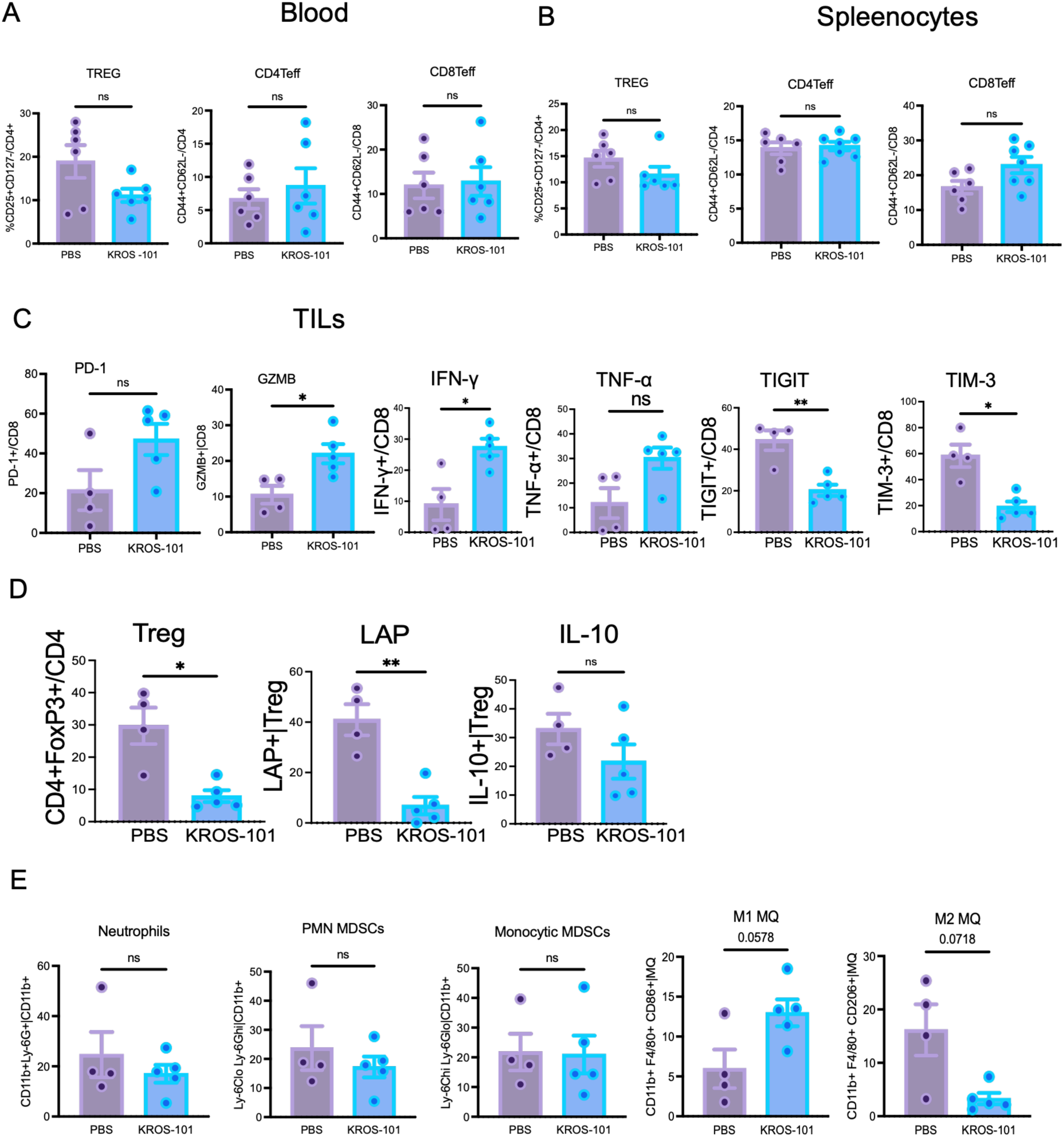
Immune profiling following KROS-101 treatment in the B16-F10 tumor model. **a**, Frequencies of Tregs (CD25+CD127− within CD4+ T cells), CD4+ effector T cells (CD44+CD62L−), and CD8+ effector T cells in peripheral blood following treatment with PBS or KROS-101. No significant differences were observed. **b**, Analysis of splenocytes showing frequencies of Tregs, CD4+ effector T cells, and CD8+ effector T cells following treatment with PBS or KROS-101. No significant differences were observed. **c**, Phenotypic and functional analysis of tumor-infiltrating CD8+ T cells. KROS-101 increased granzyme B and IFN-γ expression and reduced TIGIT and TIM-3 expression, whereas PD-1 and TNF-α were not significantly altered. **d**, Intratumoral Treg frequency and expression of the suppressive markers LAP and IL-10. KROS-101 reduced Treg frequency and LAP expression, whereas IL-10 was not significantly altered. **e**, Tumor-associated myeloid populations. KROS-101 did not significantly alter neutrophil, PMN-MDSC, or monocytic MDSC frequencies. M1-like (CD86+) macrophages showed an increasing trend, whereas M2-like (CD206+) macrophages showed a decreasing trend, neither of which reached statistical significance.

**Supplementary Fig. 5:**
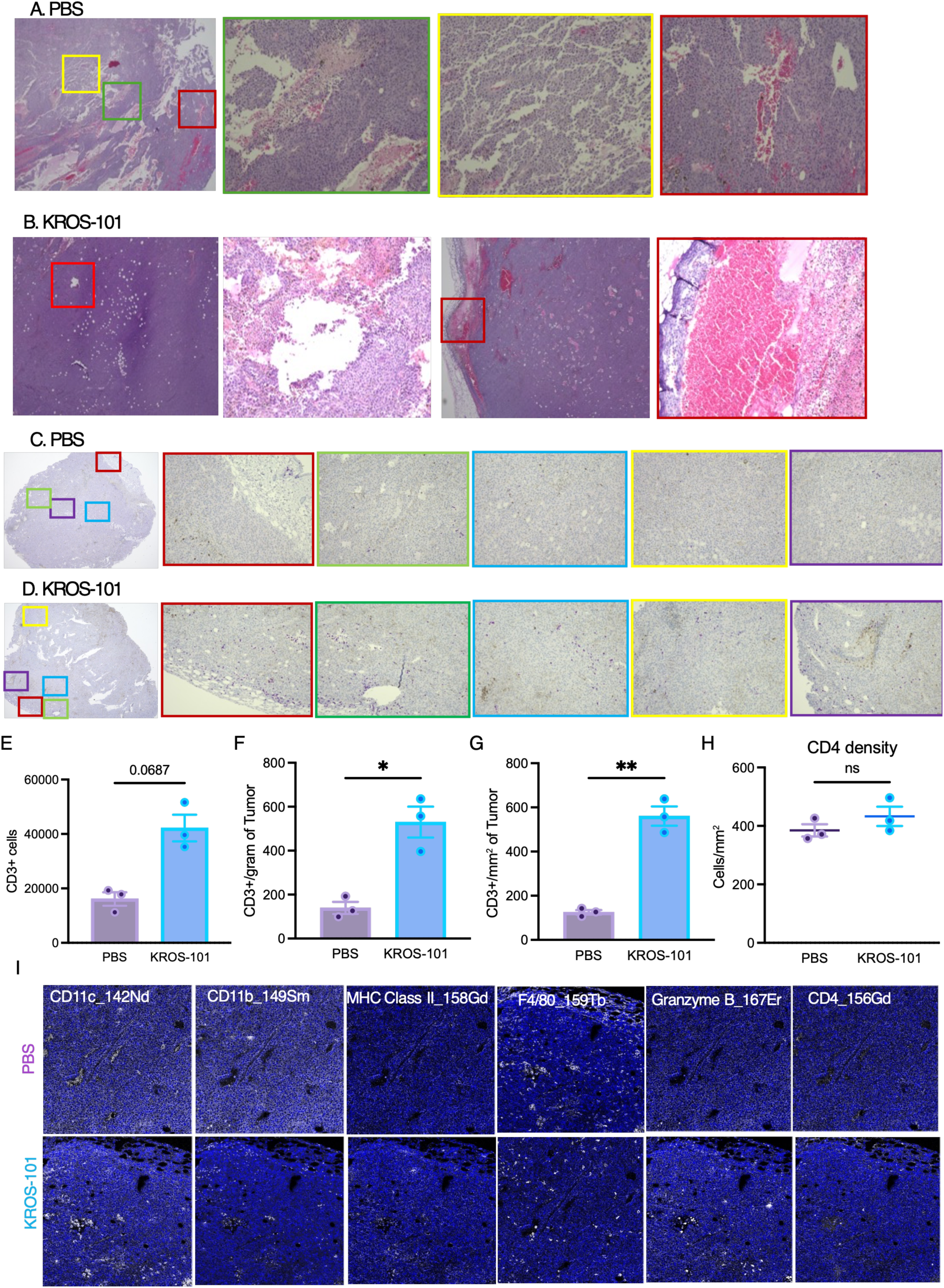
Histological and spatial immune analysis of tumors following KROS-101 treatment. **a,b**, Representative hematoxylin and eosin (H&E) stained tumor sections from PBS-treated (**a**) and KROS-101-treated (**b**) mice. Colored boxes indicate regions shown at higher magnification. **c,d**, Whole-tumor section imaging illustrating tissue architecture and CD3 T cells infiltration in PBS-treated (**c**) and KROS-101-treated (**d**) tumors with representative magnified regions highlighted. **e-g**, Quantification of tumor-infiltrating T cells measured by CD3 staining. Total CD3⁺ cells per tumor (**e**), CD3⁺ cells normalized to tumor weight (**f**), and CD3⁺ cell density per mm² of tumor tissue (**g**) show increased T cell infiltration in tumors from KROS-101-treated mice. **h**, Representative imaging mass cytometry (IMC) marker channels from tumor sections showing expression of CD11c, CD11b, MHC class II, F4/80, GZMB and CD4 in PBS-and KROS-101-treated tumors, illustrating increased immune infiltration and activation following treatment.

**Supplementary Fig. 6:**
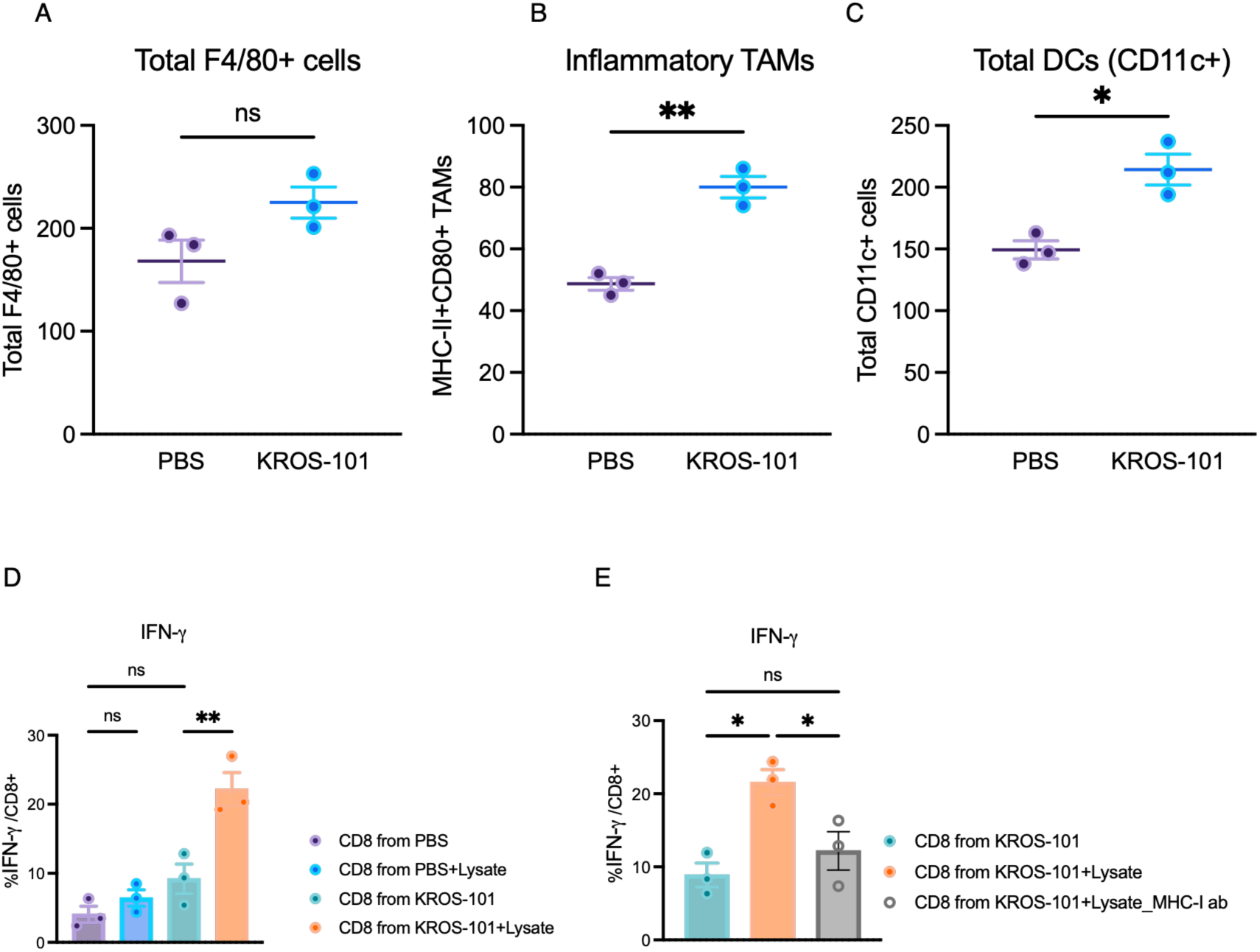
KROS-101 promotes inflammatory myeloid responses and antigen-dependent CD8⁺ T cell activation. **a**, Total F4/80⁺ tumor-associated macrophages (TAMs) in tumors from PBS- and KROS-101–treated mice, showing no significant change in overall macrophage abundance. **b**, Frequency of inflammatory TAMs (MHCII⁺CD80⁺) demonstrating increased inflammatory macrophage polarization following KROS-101 treatment. **c**, Total dendritic cells (CD11c⁺) within tumors, showing increased dendritic cell infiltration following KROS-101 treatment. **d**, IFN-γ production by CD8⁺ T cells isolated from PBS-or KROS-101–treated mice following stimulation with tumor lysate, demonstrating enhanced antigen-specific CD8⁺ T cell responses following KROS-101 treatment. **e**, IFN-γ production by CD8⁺ T cells from KROS-101-treated mice following stimulation with tumor lysate in the presence or absence of MHC class I blocking antibody, showing reduced cytokine production upon MHC blockade, indicating antigen presentation-dependent activation.

**Supplementary Fig. 7:**
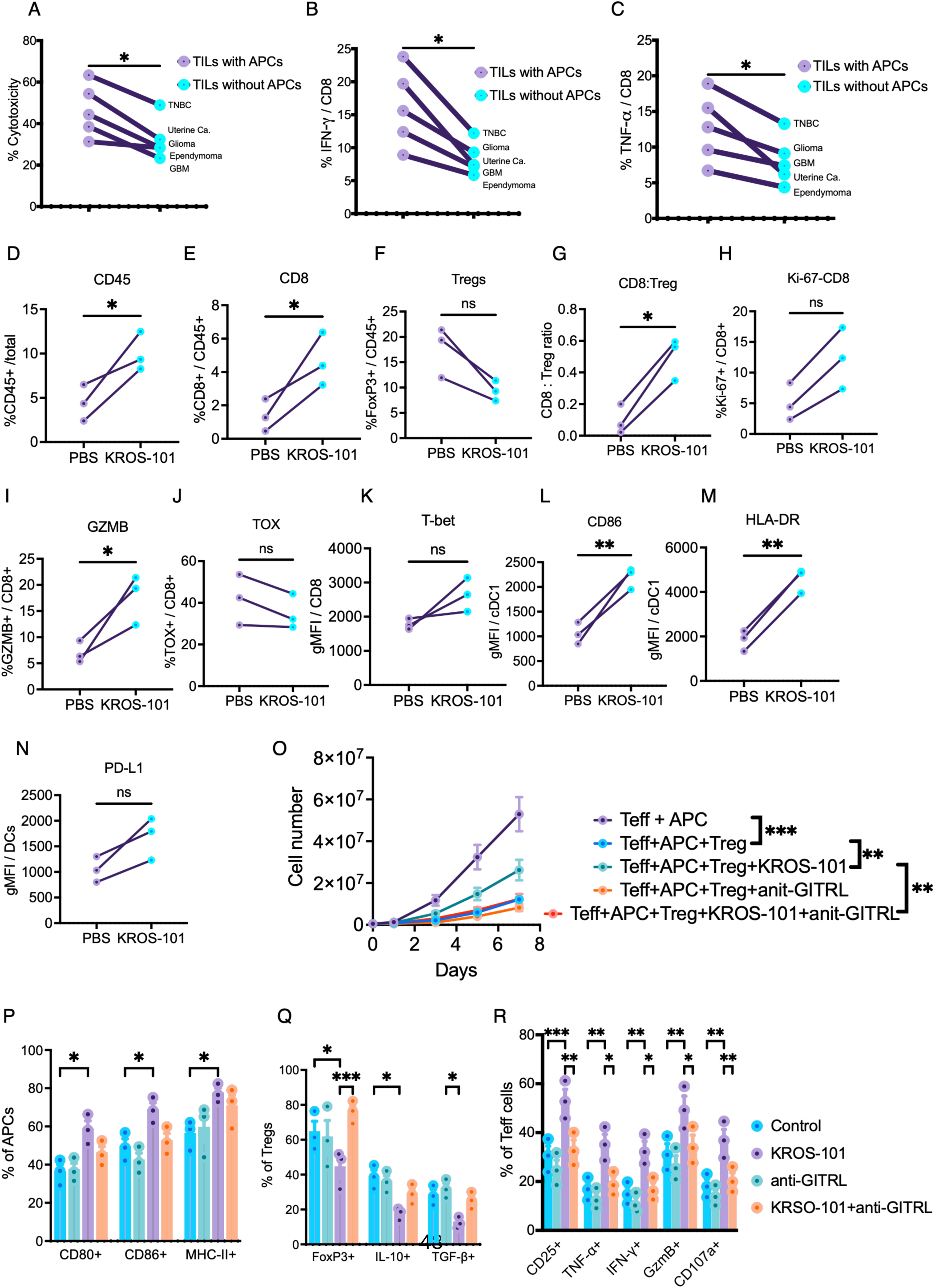
APCs enhance cytotoxic and cytokine responses of TILs and mediate the immunomodulatory effects of KROS-101. **a**, Cytotoxic activity of TILs isolated from glioma, glioblastoma (GBM), triple-negative breast cancer (TNBC) brain metastasis, ependymoma, and uterine carcinoma brain metastasis measured in the presence or absence of APCs. Co-culture with APCs significantly increased tumor cell killing. **b, c**, IFN-γ (**b**) and TNF-α (**c**) production by CD8+ TILs across tumor types in the presence or absence of APCs, showing increased cytokine responses in the presence of APCs. **D-n**, Immunophenotyping of TIL cultures isolated from glioblastoma (GBM) from three donors following treatment with PBS or KROS-101. KROS-101 increased the frequency of CD8+ cells within CD45+ cells (**d, e**) and the CD8:Treg ratio (**g**), without significantly altering Treg frequency (**f**) or CD8+ T cell Ki-67 expression (**h**). KROS-101 increased granzyme B expression (**i**), whereas TOX and T-bet were not significantly altered (**j, k**). KROS-101 also increased CD86 and HLA-DR expression on cDC1s (**l, m**), while PD-L1 expression on DCs was not significantly altered (**n**). **o**, Expansion of effector T cells (Teff) cultured with APCs in the presence or absence of Tregs, KROS-101, and/or anti-GITRL neutralizing antibody. **p**, Expression of CD80, CD86, and MHC-II on APCs following treatment with control, KROS-101, anti-GITRL neutralizing antibody, or KROS-101 plus anti-GITRL neutralizing antibody. **q**, Frequencies of FoxP3+, IL-10+, and TGF-β+ Tregs following the treatments as in (**p**). **r**, Frequencies of CD25+, TNF-α+, IFN-γ+, granzyme B+, and CD107a+ effector T cells following treatment with control, KROS-101, anti-GITRL neutralizing antibody, or KROS-101 plus anti-GITRL neutralizing antibody.

Other Supplementary Material for this manuscript includes the following:

Supplementary Table 1. Flow panel for mouse immune cells.

Supplementary Table 2. Flow panel for human immune cells.

